# Striatal and frontal signatures of social context and cost-benefit decision making in developmental stuttering

**DOI:** 10.64898/2026.03.02.707906

**Authors:** Nicole E. Neef, Elena Winter, Hellmuth Obrig, Andreas Neef, Toralf Mildner, Shaza Haj Mohamad, Christian H. Riedel, Katharina Scholze

## Abstract

Developmental stuttering is usually framed as a sensorimotor disorder, yet it manifests in communicative situations that engage motivational, self-referential, and regulatory processes. In this case-control study, we combined a socio-economic decision task outside the scanner with a socially modulated speech task during functional MRI to test how listener presence and self-referential speech shape neural activity in 34 adults who stutter and 32 controls. Both groups valued talking with another person about themselves more than talking with another person about someone else or talking to themselves. On the neural level, listener-directed (vs. private) speech and self-disclosure (vs. guessing the preferences of a famous other) elicited stronger responses in the ventral striatum and medial prefrontal cortex, with no significant group differences in the central listener-directed versus private contrast. Within the stuttering group, individual differences revealed a systematic reweighting of socially modulated activation as a function of symptom burden: higher anticipation of stuttering and greater self-reported impact of stuttering on everyday life were associated with stronger engagement of motivational circuitry, greater recruitment of frontal evaluative-control regions, and reduced contextual differentiation within speech-language cortex. Stuttering anticipation and lived experience gradually shift the balance between control, language, and motivational salience-processing systems, contributing to the disorder’s marked heterogeneity and context sensitivity. These findings indicate system-level signatures of the interaction between social context and symptom severity in developmental stuttering, extending accounts based on sensorimotor deficits.

**Graphical Abstract:** Listener presence and self-disclosure give speaking social value. In adults who stutter, the difference in activity between shared (listener-directed) and private speech was modulated by stuttering anticipation. With higher anticipation, this difference increased in the nucleus accumbens (motivational salience) and in medial prefrontal and lateral orbitofrontal cortex (evaluative control), and decreased in inferior frontal and posterior superior temporal cortex (speech-language). This suggests a reweighting of cortico-basal ganglia loops towards limbic and evaluative control circuits and away from speech-language cortex. Created with BioRender.com.

**ALT TEXT:** Graphical abstract showing private and shared speaking scenes linked by a monetary trade-off, and brain views marking accumbens and prefrontal regions with upward arrows and inferior frontal and superior temporal regions with a downward arrow.

## Introduction

Human speech is inherently social. Speaking to a listener carries motivational relevance ^1^ because it serves motivationally salient interpersonal goals such as affiliation ^2,3^, self-disclosure ^4–6^, and social evaluation ^7,8^. Listener presence fundamentally changes the incentives and consequences of speaking, as directed speech involves reputational and interpersonal stakes. By contrast, speech produced without a listener lacks these social consequences. Thus, social context modulates the value of speaking, and directed speech may recruit social-motivational control processes that shape speech production in addition to lower-level sensorimotor mechanisms.

This social-context-dependent valuation of speaking is particularly relevant for developmental stuttering, a common speech disorder characterized by repetitions of sounds and syllables, prolongations, and blocks ^9^, and by pronounced sensitivity to social-evaluative context ^10–12^. Stuttering is highly situationally variable ^13,14^ and is markedly reduced during private speech ^15,16^ when utterances are not directed to a listener or embedded in a communicative goal. By contrast, adults who stutter (AWS) frequently report increased difficulty when speaking under conditions of audience presence or communicative pressure ^17^. On the neural level, perceived discomfort is associated with right amygdala activity during stuttering occurrence ^18^. Neuroimaging research has implicated alterations within cortico-basal ganglia-thalamo-cortical circuits involved in speech timing and motor control ^19–21^ yet it remains unclear how social context interacts with reward-related systems during speaking. Given the context sensitivity of stuttering, altered valuation of directed speech may represent a mechanism contributing to symptom expression ^22,23^.

A reward-related circuitry, including the nucleus accumbens and ventral striatum, supports the valuation of actions ^24^ and is reliably engaged by social rewards ^25,26^. Beyond responses to primary reward, ventral striatum is sensitive to being observed, receiving social feedback ^27,28^, and communicating self-relevant information ^29,30^. Consistent with a role in assigning motivational salience to social acts, both adults and children who stutter show altered grey-matter morphology of the ventral striatum and nucleus accumbens compared to fluent speakers and typically developing peers ^22,31,32^. Specifically, adults who stutter show increased grey matter density in voxel-based morphometry and structural shape differences in vertex-wise shape analyses of the right ventral striatum and nucleus accumbens, while children who stutter show reduced nucleus accumbens volume compared to their peers. This pattern points to an atypical developmental trajectory of striatal circuitry in stuttering. Rather than a uniform increase or decrease, it suggests that the motivational salience system for social communication is reorganized over time when language development is accompanied by a communication disorder such as stuttering. In this context, speech directed toward a listener is likely to recruit motivational circuits differently in people who stutter than in fluent speakers, with downstream effects on speech motor and control systems in a listener dependent manner.

Given that ventral striatum encodes the motivational salience of social communication and shows atypical development in stuttering, we aimed to test whether listener presence and self-referential content specifically modulate speech-related activity within this circuitry and its interaction with language and control networks. We investigated how social context modulates neural responses during speech, and whether this modulation differs between adults who do or do not stutter (AWS/AWNS). Participants completed a monetary decision task in which they either disclosed their own preferences or inferred the preferences of a well-known person, under share conditions (speech audible to the experimenter) or private conditions (speech inaudible) ^29,30^. This design enabled us to isolate the effect of directed speech (share) versus non-directed speech (private) while controlling for speech content. From a computational perspective, our paradigm requires participants to integrate social value, internal costs (e.g. effort, anticipated dysfluency), and monetary reward, engaging multiple valuation and control systems in addition to speech motor mechanisms rather than a unitary ‘speech’ module. Small monetary incentives serve as a utility metric to reveal preference structure through real economic choices: the diagnostic value lies in the relative trade-off between monetary reward and communicative preference, rather than in the absolute incentive magnitude. This follows proposals that ecologically valid paradigms combining several value dimensions are particularly informative for understanding neurodevelopmental and mental health-related differences in social behavior ^33^. We hypothesized that directed speech would increase activity in reward-related circuitry, particularly the nucleus accumbens, and that AWS would show altered social-context-dependent modulation relative to AWNS. To relate neural responses to subjective valuation, we additionally quantified the relative monetary value of speaking in these conditions in an independent behavioral examination ^29^.

## Materials and methods

### Participants

Thirty-four adults who stutter (AWS; 29 males, five females; aged 19-46 years; one left-hander) and 32 age- and sex-matched adults who do not stutter (AWNS; 28 males, four females; aged 21-46 years; one left-hander) took part in a larger project on persistent developmental stuttering. Handedness was assessed with the Edinburgh Inventory ^34^. AWS were recruited via advertisements and social media of the German Stuttering Association; AWNS were recruited from the Max Planck Institute for Human Cognitive and Brain Sciences volunteer database. Stuttering in AWS had been previously confirmed by a speech-language pathologist, and all self-identified as people who stutter. AWNS reported no personal or family history of stuttering or other speech or language disorders. All participants had normal or corrected-to-normal vision, met MRI safety criteria, and provided written informed consent. This study was approved by the University of Leipzig ethics committee [123/17-ek].

Stuttering severity of AWS was scored using the Stuttering Severity Instrument-4 (SSI-4) ^35^ and the German Version of the Overall Assessment of the Speaker’s Experience of Stuttering (OASES) ^36,37^. SSI-4 outcomes ranged from very mild (n = 11), mild (n = 8), moderate (n = 6), severe (n = 2), to very severe (n = 7). OASES outcomes ranged from mild (n = 1), mild-to-moderate (n = 6), moderate (n = 16), moderate-to-severe (n = 7), severe (n = 2) to very severe (n = 1). Anticipation of stuttering was measured with the Premonitory Awareness in Stuttering Scale (PAiS) ^38^, a 12-item self-report questionnaire adapted from the Premonitory Urge for Tics Scale ^39^. Example items include: “Right before I stutter, I can sense that I am about to stutter,” and “Right before I stutter, I feel tense.” Trait and state anxiety were assessed with the German version of the State-Trait Anxiety Inventory ^40^, revealing modestly elevated but non-clinical anxiety levels in AWS relative to AWNS (Table 1, Supplementary Fig. 1). OASES, PAiS and State-Trait Anxiety Inventory data were not available for one AWS; analyses involving these measures were therefore based on n = 33. Professional education did not differ between groups, whereas AWNS more often had the highest school education level (Table 1, Supplementary Fig.2).

**Table 1.** Demographic details of participants, clinical features of patients.

|  | AWNS<br>(*n*=32) | AWS<br>(*n*=34) | Difference, (*P*) |
|---|---|---|---|
| Age in years, mean (SD) | 32.1 (6.7) | 31.2 (6.4) | 0.57 (0.574)<sup>b</sup> |
| Sex, male (%) | 28 (88) | 29 (85) | 0.83 (1)<sup>c</sup> |
| Handedness, *Mdn* LQ (IQR) | 100 (12.1) | 90 (21.5) | 619.5 (0.300)<sup>d</sup> |
| School education, *Mdn* (IQR) | 4 (0) | 4 (1) | **679.0 (0.033)** <sup>d</sup> |
| Professional education, *Mdn* (IQR) | 7 (2) | 6 (3) | 645.0 (0.176)<sup>d</sup> |
| Trait anxiety percentile rank, *Mdn* (IQR) | 27 (40) | 65 (49)<sup>a</sup> | **253.5 (< 0.001)** <sup>d</sup> |
| State anxiety percentile rank, *Mdn* (IQR) | 16 (43) | 51 (38)<sup>a</sup> | **318.0 (0.005)** <sup>d</sup> |
| SSI-4 overall score, *Mdn* (IQR) | 0 (0) | 22 (14.5) | **0 (< 0.001)** <sup>d</sup> |
| % stuttered syllables, *Mdn* (IQR) | 0 (0.3) | 5.5 (8.8) | **0 (< 0.001)** <sup>d</sup> |
| OASES total score, *Mdn* (IQR) | n/a | 2.8 (0.8)<sup>a</sup> | n/a |
| PAiS, *Mdn* (IQR) | n/a | 18 (7.8)<sup>a</sup> | n/a |

LQ = laterality quotient; IQR = interquartile range; SSI = Stuttering Severity Index;(35) OASES = Overall Assessment of the Speaker´s Experience of Stuttering; (37,41) Premonitory awareness in stuttering;(38); school and professional education were coded as ordinal variables. Highest school qualification was classified as: (0) no qualification, (1) general education school-leaving certificate after grade 9, (2) general education school-leaving certificate after grade 10, (3) university of applied sciences entrance qualification, and (4) higher education entrance qualification. Highest professional qualification was classified as: (0) no qualification; (1) vocational training <1 year; (2) vocational training <2 years; (3) vocational training <3 years; (4) completed vocational qualification; (5) currently enrolled at a university or university of applied sciences; (6) bachelor’s degree or master craftsman qualification; (7) master’s degree or equivalent; and (8) doctoral degree or higher. an = 33; differences were tested with unpaired, two-tailed t-test (R command: t.test), cFisher’s Exact Test (R command: fisher.test), and d Wilcoxon Signed-Rank Test (R command: wilcox.exact).

### Behavioural task: subjective value of speaking

The behavioural paradigm was adapted from Tamir and Mitchell ^29^ to assess social decision-making under monetary trade-offs. Participants chose between answering a question aloud or waiting passively, allowing quantification of the subjective value assigned to speaking in different social contexts. Four conditions were tested*: self share* (verbal report of one’s own preference to a listener), *self private* (spoken but unheard), *other share* (verbal report of beliefs about a public figure’s preferences to a listener), and *other private* (spoken but unheard). Angela Merkel was selected as the “other” due to her high familiarity at the time of the study, enabling plausible belief formation. Each trial required a two-alternative forced choice between speaking and waiting for 5 seconds. Both options were associated with small monetary outcomes (€0.01 - €0.04), with the relative payoff between speaking and waiting varying randomly from −€0.03 to +€0.03. In waiting trials, participants viewed a fixation cross and performed no task. Participants were seated in a sound-attenuated booth, while the experimenter (listener) was located in a separate room (Fig. 1A). Trial type, options, and payoffs were displayed at trial onset. Choices were made via button press. When speaking was selected, participants read the question aloud and responded “yes” or “no.” Example questions included *“Mag ich Flamingos?”* (*Do I like flamingos?*) in a *self*-condition trial and *“Mag Merkel Tango?”* (*Does Merkel like Tango?)* in an *other-*condition trial. The stimuli used in the behavioural experiment are listed in the Supplementary Material. Speech was recorded in all conditions but transmitted to the listener only during share trials. Prior to the experiment, practice trials demonstrated the share versus private manipulation; participants wore the listener’s headphones in which the instructor gave spoken responses across all conditions, allowing participants to directly experience what the listener could and could not hear. The task modified the original button-press paradigm by requiring overt speech, permitting assessment of speech production alongside socio-economic decision-making. Each condition was presented 60 times (240 trials total), randomly distributed across four runs (7:30 min each). Item assignment and trial order were randomized per participant. The task duration was approximately 45 minutes. Stimulus presentation, recordings of the utterances and playback to the listener were controlled using Presentation software (Neurobehavioral Systems). Questions concerned everyday, non-controversial preferences.

**Figure 1.**
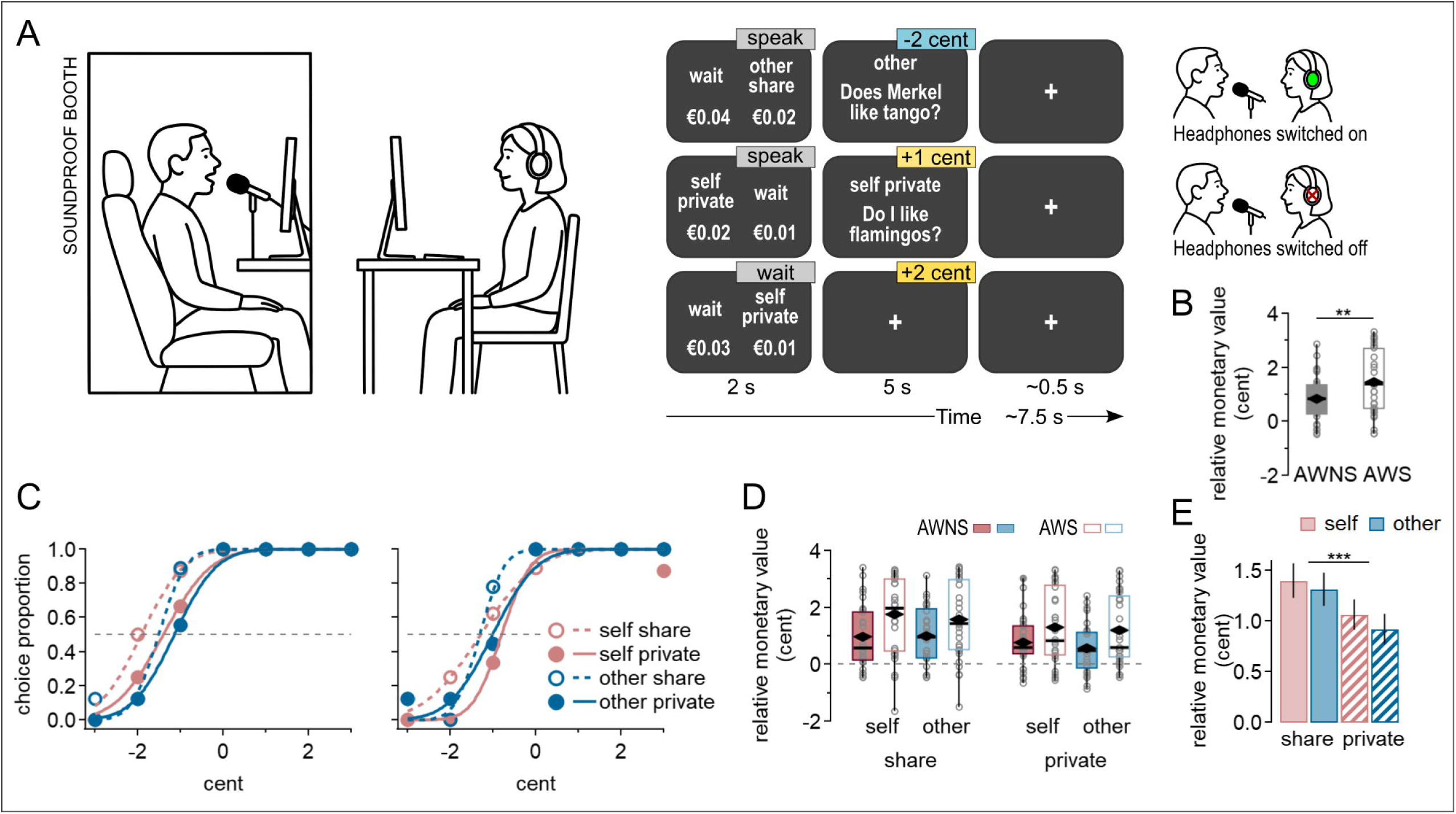
Task outside the MRI scanner and results. (A) Experimental setup. Participants were seated in a soundproof booth and viewed the stimuli on a screen. The investigator, located outside the booth, viewed the same stimuli and listened to participants’ responses during share trials only. Example trials were labeled with response options (grey boxes) and their associated monetary outcomes (gain: yellow box, loss: blue box). (B) Relative monetary values for AWNS (*n* = 27) and AWS (*n* = 32). Relative monetary value is the sign-inverted PSE. Asterisks indicate a significant main effect of Group in a robust linear mixed model controlling for trait anxiety and professional education (*t*(55) = -2.90, *p* = .005); ** *p* < .01. (C) Representative individual data showing PSEs from one AWNS (left) and one AWS (right). The AWNS example shows the most negative PSE in the self-share condition, indicating a willingness to forgo 2 cents to share self-related information. The AWS example shows negative PSEs for both self- and other-related conditions, indicating a willingness to forgo approximately 1.5 cents to speak. (D) Box plots show the median relative monetary values (- PSEs) and interquartile range, with whiskers indicating the minimum and maximum and diamonds indicating the mean value for AWNS (*n* = 27) and AWS (*n* = 32). For (B) and (D), values are sign- flipped relative to the fit curve; positive values reflect willingness to pay to speak (E) Mean relative monetary value (-PSEs) for each of the four conditions, pooled across all participants. Each bar is based on *n* = 59 participants (AWNS: *n* = 27, AWS: *n* = 32, one value per participant and condition); error bars indicate the standard error of the mean (SEM). Asterisks indicate a significant main effect of Mode (share vs private) in the robust linear mixed model (*t*(171) = -4.56, p < .001). *** p < .001. **ALT TEXT:** Panel A shows the laboratory setup and example trials for each condition of the behavioral socioeconomic decision task. Panels B to E show the results: higher relative monetary values in adults who stutter than in adults who do not stutter (B), example choice curves from one participant per group (C), relative monetary values for each condition and group (D), and higher values for share than for private conditions (E).

**Figure 2.**
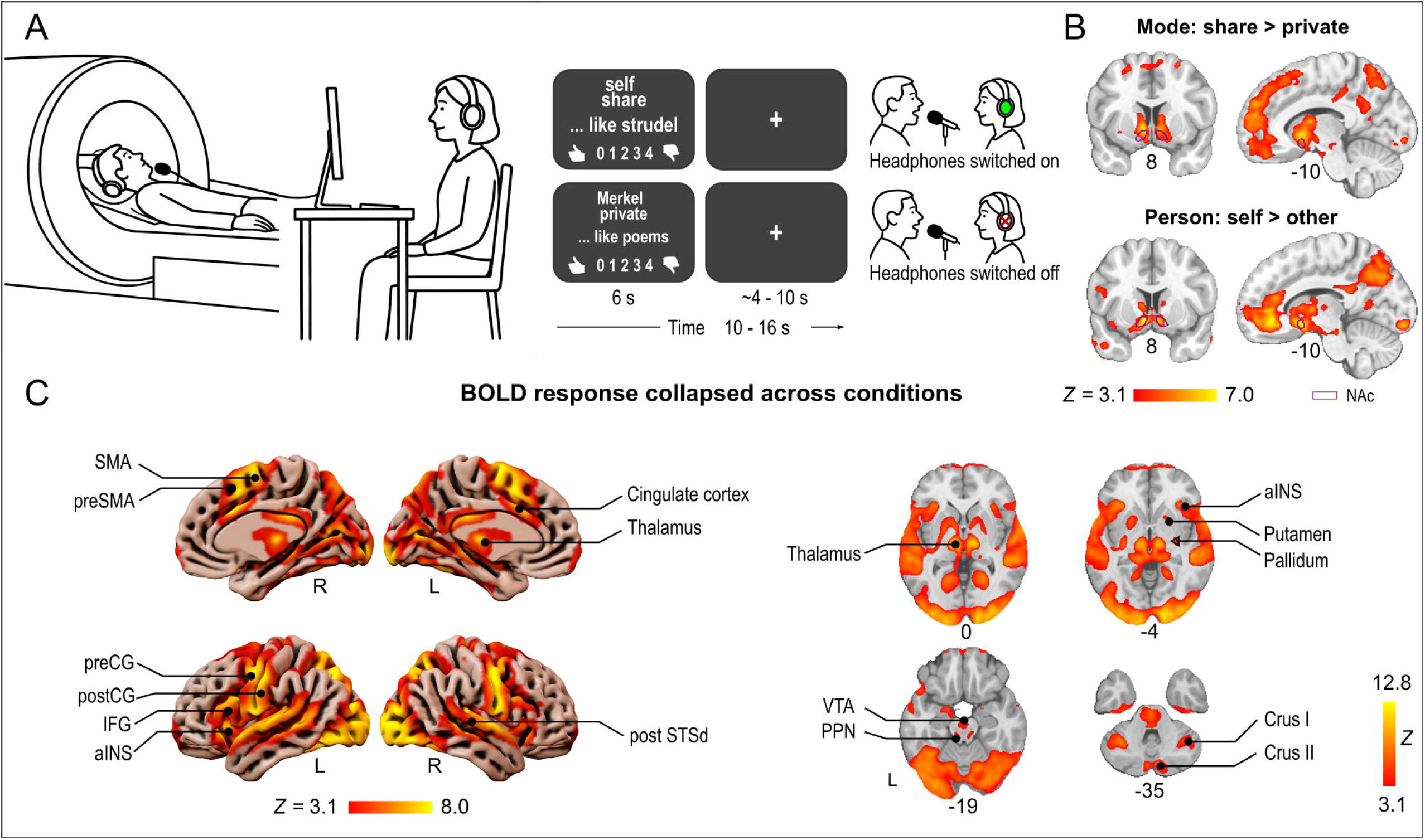
Task inside the MRI scanner and results pooled across groups. (A) Experimental setup. Participants lay in the MRI scanner and viewed the stimuli and responded to each trial. The investigator, located outside the scanner room, viewed the same stimuli and listened to participants’ responses during share trials only. (B) Whole-brain analysis revealed a main effect of Mode, with greater nucleus accumbens (NAc) responses during share compared with private trials, and a main effect of Person, with greater NAc responses during self compared with other trials (results pooled across groups, *n* = 66). (C) For visualization purposes, we performed a whole-brain analysis across all conditions (*n* = 66) to illustrate task-positive speech-related activity. All fMRI results are based on FLAME (FMRIB’s Local Analysis of Mixed Effects) stage 1. Contrast maps are cluster-corrected using a cluster-forming threshold of *Z* > 3.1 and *p* < .05. Abbreviations: aINS, anterior insula; IFG, inferior frontal gyrus; NAc, Nucleus accumbens; PPN, pedunculopontine nucleus; postCG, postcentral gyrus; post STSd, dorsal posterior superior temporal gyrus; preCG, precentral gyrus; preSMA, presupplementary motor area; SMA, supplementary motor area; VTA, ventral tegmental area. **ALT TEXT:** Panel A shows the setup and examples of the conditions of the task during scanning. Panels B and C show the results: greater nucleus accumbens activity for share than for private and for self than for other trials (B), and widespread speech-related brain activity across all conditions (C).

### Statistical analysis of behavioural data

To estimate the subjective value of speaking, we computed the proportion of trials in which the speaking option was chosen at seven relative payoff levels (-€0.03 to +€0.03) for each participant and condition. Following Tamir & Mitchell (2012), small monetary incentives were used deliberately as a utility metric to elicit real economic choices, rather than to represent the absolute psychosocial cost of stuttering. The diagnostic value of this approach lies in choice behaviour, specifically, the relative trade-off between monetary reward and communicative preference, rather than in the absolute incentive magnitude. These data were used to estimate the point of subjective equality (PSE), defined as the payoff difference at which speaking and waiting were chosen with equal probability, thereby indexing the subjective value of speaking relative to silence. Choice probabilities were fitted with cumulative normal functions using a Levenberg-Marquardt least-squares algorithm (Igor Pro 9, Wavemetrics, Fig. 1C). Negative PSE values indicate a willingness to incur a monetary cost to speak, whereas positive values indicate that speaking required financial compensation. In cases where choice behaviour provided insufficient constraint (e.g. near-deterministic responding), weakly informative Gaussian priors were applied to stabilise parameter estimation without materially affecting well-constrained fits. Where fits remained unconstrained despite priors (near-exclusive choice of speaking; AWS: n = 2/34; AWNS: n = 5/32), these participants were excluded, leaving n = 32 AWS and n = 27 AWNS for PSE group analyses.

Sign-flipped PSE values (such that positive values reflect willingness to pay to speak) were analyzed using a robust linear mixed model (*rlmer*) ^41^ with the between-subjects factor Group (AWNS versus AWS) and the within-subjects factors Mode (share versus private) and Person (self versus other), and trait anxiety and professional education included as nuisance covariates. Subject was modeled as a random intercept to account for repeated measures. Factors were sum-coded. *P*-values were derived using Satterthwaite-approximated degrees of freedom from the corresponding non-robust model (*lmerTest*). Group statistics were calculated with RStudio 2022.07.1 using the *robustlmm* library.

### Determination of stuttering in experimental recordings

Recordings from the behavioural experiment were assessed for stuttering occurrence with a scoring system (0 for no stuttering, 1 for stuttering). Speech blocks, prolongations, and part-word repetitions were considered stuttering. In addition, dysrhythmic repetitions of monosyllabic words were also classified as stuttering. Speech blocks were considered to occur when an unnatural pause appeared that was longer than 300 ms and did not correspond to the expected prosodic patterns, for example, a syntactically or semantically inappropriate pause in a position where no prosodic boundary would be typically expected. Pauses were also considered stuttering in case an audible interruption (glottis noise) occurred in the otherwise continuous phonation. To ensure conservative and reliable coding, ambiguous instances or unclear audio recordings were not labelled as stuttering. Other types of disfluencies, such as polysyllabic whole-word repetitions and interjections (e.g., “um,” “like”), were not considered stuttering. Stuttering response rate was then calculated by dividing the number of stuttered trials by the total number of share trials for each participant. We used paired Wilcoxon Signed-Rank tests to assess whether the condition speaking privately or to someone else influenced the stuttering response rate in individuals who stutter.

To assess inter-rater reliability, a second rater independently evaluated 10% of the trials. Participant selection was stratified to reflect the overall distribution of stuttering severity in the full sample, resulting in a subset that included one participant with mild stuttering, six with moderate stuttering, two with mild-to-moderate stuttering, two with moderate-to-severe stuttering, and one with severe stuttering. For each of these 12 participants, the spoken trials from one run were analysed, with one of the four runs per participant chosen pseudo-randomly. This yielded a total of 549 reanalysed trials.

Cohen’s Kappa was calculated to estimate inter-rater reliability, accounting for the binary nature of the data and its imbalanced distribution. The analysis yielded a Cohen’s Kappa of 0.81 (95% CI [ 0.75, 0.87], indicating strong agreement according to McHugh, 2012 ^42^.

As task familiarity or habituation may influence stuttering behaviour over time, we conducted a Friedman test to examine whether the proportion of stuttered trials varied across runs. The analysis revealed no significant effect of time, *X*²(3) = 0.97, p = 0.81, indicating that the proportion of stuttered trials remained stable throughout the experiment.

### fMRI task: social modulation of speech

The fMRI paradigm was designed to assess whether social context modulates nucleus accumbens activity. To maximise the number of active trials, the waiting option used in the behavioural task was removed. Participants completed the same four conditions (*self share*, *self private*, *other share*, *other private*) and responded on every trial using a 5-point Likert scale ranging from *extremely unlikely* to *extremely likely.* In the self-conditions, the scale indicated the extent to which the statement reflected their own preferences; in the other conditions, it indicated how likely they believed the statement was to reflect Angela Merkel’s preferences. Participants indicated their response verbally by stating a number between 0 and 4. Unlike in the behavioural experiment, they did not read the statements aloud. During share trials, a study assistant outside the scanner listened to participants’ responses via headphones (Fig. 2A).

Example statements included “Ich mag Strudel.” (I like strudel.) in a self-condition trial and “Merkel mag Gedichte.” (Merkel likes poems.) in an other-condition trial. The stimuli used in the fMRI experiment are listed in the Supplementary Material. Each condition was presented 48 times (192 trials total), randomly distributed across four runs in an event-related design (9:40 min each). Trial order and item assignment were randomized per participant. The task lasted approximately 45 minutes and was implemented using Presentation software (Neurobehavioral Systems). All stimuli were novel and did not overlap with those used in the behavioural experiment. Complete lists of stimuli are provided in the Supplementary Material.

Participants’ verbal responses were recorded using a fibre-optic, dual-channel, noise-cancelling microphone (FOMRI III, OptoAcoustics Ltd., Israel) attached to the head coil. The signal was routed through an MR Confon amplifier (MR Confon GmbH, Magdeburg, Germany), filtered to attenuate scanner noise, and transmitted to the sound card of the stimulus presentation computer. Trial-level playback was controlled via Presentation software (Neurobehavioral Systems, Inc.), which routed the audio signal to the listener’s headphones during share trials and suppressed playback during private trials. Training was conducted outside of active scanning; participants therefore experienced the share/private manipulation in the presence of scanner background noise (cooling pumps) but not the acoustic environment of active MRI acquisition. During scanning, the operator’s audio feed was available throughout the session for safety monitoring and was used to verify correct signal acquisition at the start of the experiment. Continuous safety monitoring was ensured through standard MRI safety procedures: participants’ cardiac pulse and oxygen saturation were monitored via pulse oximetry throughout the session, and participants held an alarm ball with explicit instructions to use it should they feel uncomfortable or wish to stop at any point.

Stuttering was not accessed from the fMRI recordings, as the combination of continuous scanner noise and monosyllabic verbal responses (digits 0–4) precluded reliable classification.

### MRI data acquisition

Whole brain fMRI data were acquired continuously using a 3 Tesla MAGNETOM Prismafit scanner (Siemens Healthineers, Erlangen, Germany) equipped with a 32-channel receive head coil and a dual gradient-echo echo planar imaging sequence.^43^ Scanning parameters were as follows: repetition time (TR) = 2000 ms; echo times (TEs) = 12/33 ms; partial Fourier factor = 6/8; bandwidth = 1966 Hz/Px; flip angle = 90°; voxel size = 2.5 × 2.5 × 2.5 mm³; interslice gap = 0.25 mm; and field of view (FOV) = 204 mm. Image readout was accelerated by using GeneRalized Autocalibrating Partial Parallel Acquisition (GRAPPA) and multiband (SMS) acquisition, both by a factor of 2. Across 4 runs, a total of 1160 volumes were acquired per participant, each consisting of 60 slices aligned to the AC–PC plane and acquired in interleaved order. Before the start of the stimulation paradigm in each run, 30 auxiliary volumes were taken, which were used to calculate the weighting factors for the echo combination. Additionally, sagittal-oriented T1-weighted images were acquired using a 3D magnetization-prepared 2 rapid gradient echo (MP2RAGE)^44^ sequence with the following parameters: inversion times (TIs) = 700/2500 ms; TR = 5000 ms; TE = 1.96 ms; bandwidth = 240 Hz/Px, flip angles = 4/5°, voxel size = 1 × 1 × 1 mm^3^; acquisition matrix 256 × 256 × 176; GRAPPA factor = 3; and acquisition time = 8:22 min.

### MRI data analyses

A hybrid time series consisting of a weighted sum of both individual echo time series was created using Statistical Parameteric Mapping (SPM12, Welcome Trust Centre for Neuroimaging, UCL, London, UK) and an additional script both implemented in MATLAB (The MathWorks, Natick, MA, USA) and available at: https://doi.org/10.25625/LZJNSZ. In the first step, both echo time series (including their auxiliary volumes) were re-sliced using identical alignment parameters derived from the first echo’s time series. Subsequently, the auxiliary volumes of both echoes were used to calculate maps of the respective tSNR and weighting factors. Using these weighting factor maps, the final hybrid time series for the functional volumes was created by weighted summation.^45^ Functional hybrid data were further analysed with FSL v6.0.7.16 using the FEAT pipeline ^46,47^. Preprocessing comprised spatial smoothing with a 5 mm full-width at half-maximum (FWHM) Gaussian kernel, high-pass temporal filtering (cut-off 90 s), co-registration to the individual anatomical image, and normalization to the MNI ICBM152 nonlinear asymmetric template (mni_icbm152_t1_tal_nlin_asym_09a). T1-weighted images were segmented using FastSurfer ^48^.

Head motion was quantified using framewise displacement, calculated as the sum of absolute scan-to-scan displacements across the six rigid-body realignment parameters, with rotational parameters converted to millimeters assuming a 50 mm sphere radius ^49^. Mean framewise displacement was low in both groups (AWS: *M* = 0.16 mm, *SD* = 0.06 mm; controls: *M* = 0.17 mm, *SD* = 0.05 mm) and did not differ significantly between groups (*t*(63.6) = -0.481, *p* = 0.632), indicating that head motion was unlikely to confound group-level results. No participants were excluded based on motion.

At the first level, a general linear model was specified with the four tasks convolved with the hemodynamic response function; temporal derivatives were included as nuisance regressors. Within each participant, conditions were then combined using a fixed-effects analysis to model a 2 × 2 ANOVA with the factors Mode (share vs private), Person (self vs other), and their interaction. Contrast estimates (COPE images) were then entered into higher-level random-effects analyses to test main effects of Mode and Person, the Mode × Person interaction, and group differences (AWNS vs AWS) for each effect. To control for potential confounds, supplementary analyses reran the higher-level models including trait anxiety (STAI percentile rank) and professional education as demeaned nuisance covariates. Statistical inference was performed on the whole brain, using cluster-wise correction with a cluster-forming threshold of Z > 3.1 and a family-wise error-corrected cluster-extent threshold of p < 0.05 ^50^.

### Correlation analyses with clinical measures

We examined whether individual differences in (i) anticipation of stuttering and (ii) overall stuttering impact were associated with the modulation of brain activity by social context. To this end, we performed additional cluster-level correlation analyses testing positive and negative associations between PAiS and OASES scores and neural modulation by Mode (share > private) as well as by Person (self > other). PAiS and OASES scores were not significantly correlated (Spearman r(31) = 0.27, p = .131, two-tailed), whereas OASES scores showed a trend toward a positive association with SSI-4 total scores (Spearman r(31) = 0.31, p = .081, two-tailed). Because OASES scores showed a trend toward a positive association with SSI-4 total scores (Spearman r(31) = 0.31, p = .081, two-tailed), and to avoid redundancy and inflation of type-I error from including partially overlapping clinical measures, SSI-4 was not entered as an additional predictor in the voxel-wise correlation analyses. Instead, we focused on PAiS and OASES as conceptually distinct dimensions of stuttering experience. Age was included as a nuisance covariate. Correlation analyses were restricted to voxels within the significant clusters of the respective contrasts, and statistical inference was based on cluster-wise correction with a cluster-forming threshold of Z > 3.1 and a family-wise error-corrected cluster-extent threshold of p < 0.05. Correlation analyses were implemented in FEAT using a fixed-effects model. Given that developmental stuttering is not a homogeneous disorder^13^, together with the marked inter individual variability in neural activation patterns^23^, and the absence of robust group level differences in the random effects analyses in the critical contrast (share > private), we focused on within sample correlations between contrast estimates and clinical scores. A fixed effects model was therefore chosen to maximize sensitivity to these individual difference associations, with the understanding that inference is confined to this sample. We included age as a covariate of no interest to account for age related variance in brain activation and clinical scores, ensuring that observed associations primarily reflect stuttering related individual differences.

## Results

### Effects of group and social context on the subjective value of speaking

Statistics of the PSE values indicated a significantly higher relative monetary value of speaking in AWS compared with AWNS (Fig. 1B). A robust linear mixed model controlling for trait anxiety and professional education revealed a main effect of Group (*t* = -2.90, *p* < .005), with higher PSE values in AWS (mean = 1.47 cents, 95% CI [1.23, 1.70]) than in AWNS (mean = 0.83 cents, 95% CI [0.64, 1.03]). Trait anxiety emerged as a significant predictor (*t* = 2.01, *p* = 0.049), confirming that individual differences in anxiety contributed to PSE values independently of group membership. The model further revealed a main effect of Mode (*t = -4.56*, *p* < 0.001). Across groups, participants were willing to sacrifice monetary reward to share information rather than keep it private (shared: mean = 1.36 cents, 95% CI [1.13, 1.60]; private: mean = 0.99 cents, 95% CI [0.78, 1.20]; Fig. 1E). Neither the main effect of person (*t* = -1.42, *p* = 0.158), professional education (*t* = 1.47, *p* = 0.147), nor any of the two- or three-way interactions reached significance (all *p* > 0.33).

To further characterize the three-way interaction, we conducted post hoc unpaired *t*-tests. These analyses suggested that AWS assigned higher value than AWNS to sharing information about themselves and to speaking about others in private; however, neither comparison survived correction for multiple testing. No group differences were observed for the remaining conditions (*other share* and *self private*; Supplementary Table 1).

### No effect of listener presence on stuttering rate

We next tested whether stuttering rate, assessed from speech recorded outside the scanner during the behavioral experiment, was modulated by listener presence. In the *share* conditions, participants were aware that the experimenter could hear their speech. Conversely, in the *private* conditions they knew that their utterances were not audible to the experimenter. Stuttering rates were highly similar across share and private conditions (Supplementary Fig. 3A) and showed a strong positive relationship, such that participants who stuttered more in the share condition also stuttered more in the private condition. Stuttering rates varied markedly across individuals, consistent with the broad range of symptom severity captured by the SSI-4 (Supplementary Fig. 3B). In AWS, Wilcoxon signed-rank tests indicated no significant differences in stuttering rate between the *self share* and *self private* conditions (*V* = 308, *p* = 0.052, uncorrected), nor between the *other share* and *other private* conditions (*V* = 200, *p* = 0.153, uncorrected). Descriptive stuttering rate data and corrected p-values for both AWS and AWNS are provided in Supplementary Table 2.

### Effects of listener presence and self-relatedness on brain activity

Whole-brain analysis revealed significant effects of listener presence and self-relatedness of speech content. Speech produced in the presence of a listener (share > private) engaged widespread cortical and subcortical brain regions spanning medial and lateral prefrontal, premotor, posteromedial parietal, posterior cingulate, and temporo-parietal cortex, together with the caudate nucleus extending into the nucleus accumbens and cerebellar Crus I (Fig. 2B; Supplementary Fig. 4, Supplementary Table 3). Speech referring to the self rather than another person (self > other) recruited a distributed set of frontal and parietal association regions, including anterior cingulate, inferior frontal, and ventrolateral/orbitofrontal prefrontal cortex extending into the caudate nucleus, nucleus accumbens and thalamus, as well as precuneus and inferior parietal cortex, with additional involvement of lateral temporal and occipital regions, cerebellar Crus II, and hippocampus (Fig. 2B; Supplementary Fig. 4, Supplementary Table 3).

### Listener-related modulation of brain activity by group

Listener-related modulation of brain activity (share > private) was observed in both groups and showed a largely overlapping spatial pattern. In AWNS and AWS, directed speech engaged medial prefrontal and premotor cortex, posterior cingulate and posteromedial parietal regions, as well as subcortical structures including the caudate nucleus, thalamus, and cerebellum. Additional frontal and temporal regions and extended activity of the nucleus accumbens were evident in the AWS map (Fig. 3A), whereas medial frontal and parietal components were more prominent in AWNS (Fig. 3B); however, the direct between-group comparison was not significant (Supplementary Table 4). These activation patterns were not materially affected by trait anxiety or professional education, with the exception of thalamic modulation in AWNS, which was attenuated after covariate adjustment (Supplementary Fig. 5).

**Figure 3.**
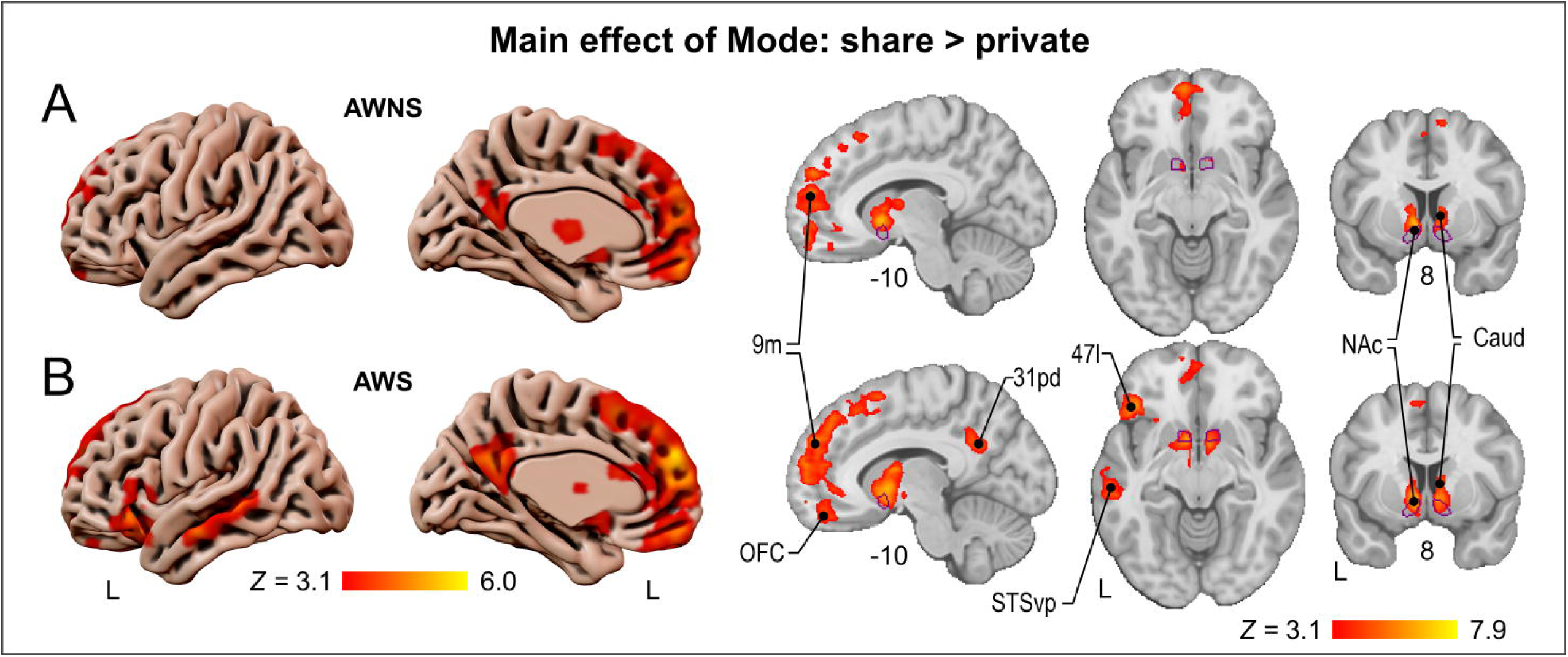
Listener presence. For the share > private contrast, both AWNS (*n* = 32) (A) and AWS (*n* = 34) (B) showed increased BOLD responses in the nucleus accumbens (NAc) and caudate nucleus (Caud), together with bilateral anterior cingulate cortex (area 9m) and thalamic activity. In AWS, additional effects were observed in the left inferior frontal gyrus, pars orbitalis (area 47l), the left ventral posterior superior temporal sulcus (STSvp) extending into middle temporal gyrus, and posterior cingulate cortex (area 31pd). All fMRI results are based on FLAME (FMRIB’s Local Analysis of Mixed Effects) stage 1. Contrast maps are cluster-corrected using a cluster-forming threshold of *Z* > 3.1 and *p* < .05. **ALT TEXT:** Panels A and B show brain surface views and sections with greater activity for share than for private trials in adults who do not stutter (A) and adults who stutter (B). Both groups show activity in the nucleus accumbens, caudate nucleus and medial frontal cortex; adults who stutter show more extensive activity, including additional left frontal and temporal regions and the posterior cingulate cortex.

### Group differences in self-related speech responses

Self-related speech (self > other) engaged frontal and parietal association cortex in both groups, with prominent involvement of anterior cingulate and ventral frontal regions. In AWNS, self-related responses additionally extended into orbitofrontal and inferior frontal cortex, inferior temporal cortex, hippocampus, and brainstem regions encompassing the ventral tegmental area and dorsal raphe nucleus (Fig. 4A). In AWS, effects were more circumscribed, centered on anterior cingulate and frontal pole regions, with additional cerebellar involvement (Fig. 4B; Supplementary Table 5). These activation patterns were not materially affected by trait anxiety or professional education (Supplementary Fig. 6).

**Figure 4.**
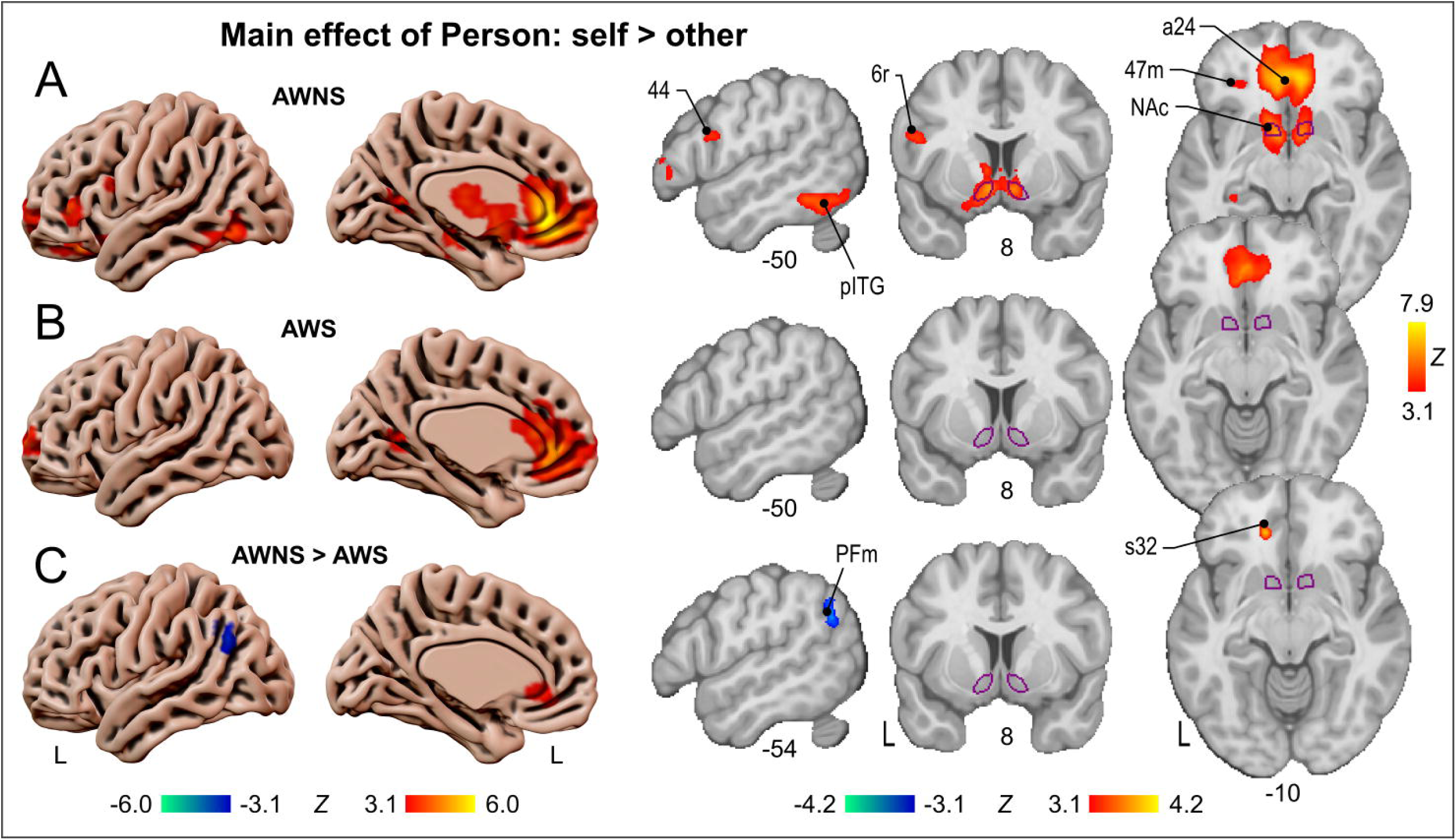
Self-related speech. For the self > other contrast, both AWNS (*n* = 32) (A) and AWS (*n* = 34) (B) showed increased BOLD responses in anterior cingulate cortex (a24) and frontal pole. In AWNS, effects extended into orbitofrontal cortex (47m) and nucleus accumbens (NAc), with additional clusters in inferior frontal gyrus (44), inferior frontal junction (6r), and posterior inferior temporal gyrus (pITG). Group differences (C) were observed in right subcallosal cortex (s32) and left inferior parietal cortex (area PFm). All fMRI results are based on FLAME (FMRIB’s Local Analysis of Mixed Effects) stage 1. Contrast maps are cluster-corrected using a cluster-forming threshold of *Z* > 3.1 and *p* < .05. **ALT TEXT:** Panels A and B show brain surface views and sections with greater activity for self than for other trials in adults who do not stutter (A) and adults who stutter (B). Both groups show activity in the anterior cingulate cortex and frontal pole; adults who do not stutter show more extensive activity, including the nucleus accumbens and additional frontal and temporal regions. Panel C shows group differences, with greater activity in adults who do not stutter in the subcallosal cortex and greater activity in adults who stutter in the left inferior parietal cortex.

Direct group comparison showed stronger self-related responses in AWNS than AWS in anterior cingulate cortex, whereas AWS showed greater responses than AWNS in medial inferior parietal cortex (Fig. 4C). The latter effect did not survive covariate adjustment for trait anxiety and professional education, suggesting that the IPL response to referential context in AWS is partially explained by individual differences in these background variables (Supplementary Fig. 6).

### Correlates of stuttering anticipation in listener-related modulation

Within AWS, stuttering anticipation (PAiS) was associated with listener-related modulation of brain activity (share > private). Higher PAiS scores correlated positively with share-related responses in the nucleus accumbens (Fig. 5A; Supplementary Table 6). In contrast, negative correlations were observed in left inferior frontal cortex, encompassing pars opercularis, triangularis, and orbitalis, as well as posterior superior temporal regions including ventral posterior superior temporal sulcus and adjacent temporal cortex (Fig. 5B; Supplementary Table 6).

**Figure 5.**
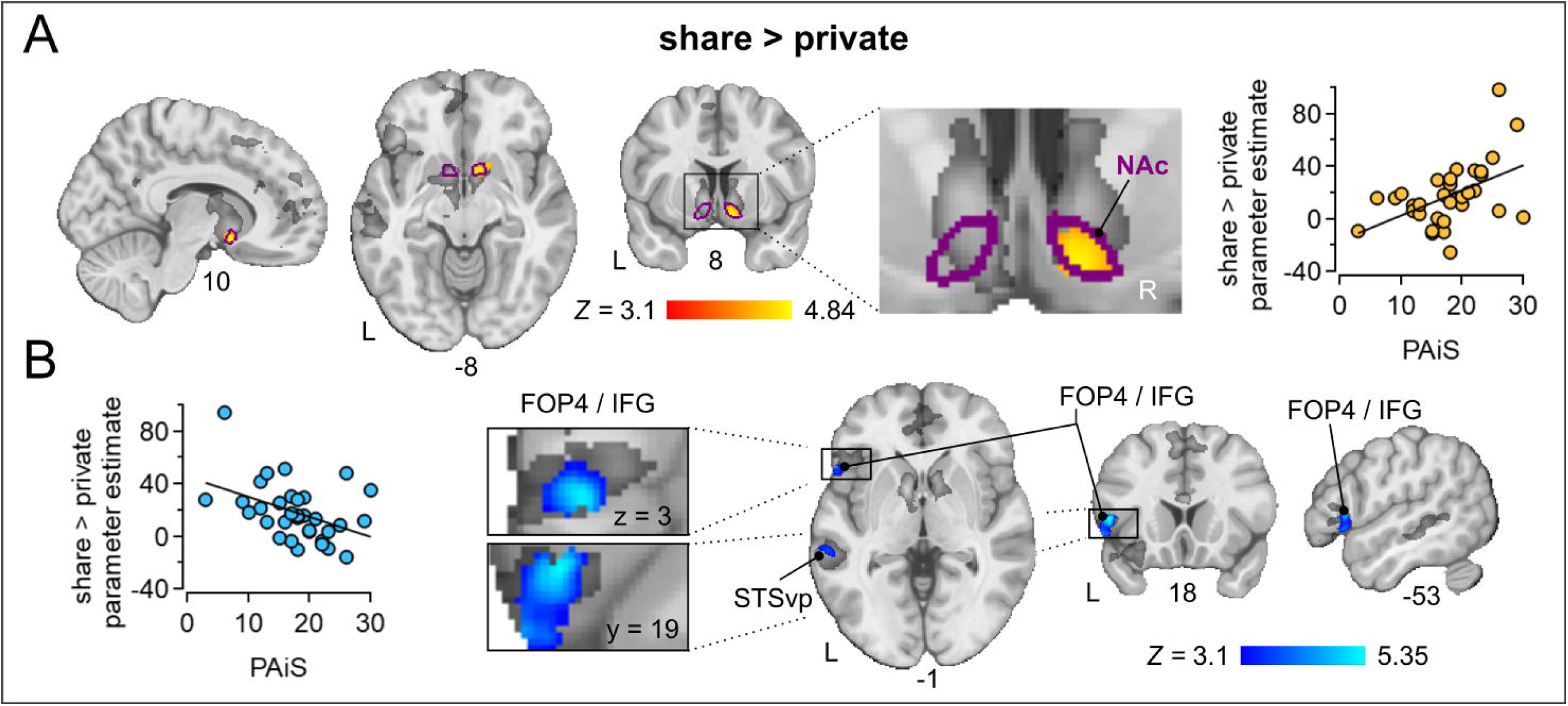
Correlates of stuttering anticipation. (A) Positive association between PAiS score and listener-related neural modulation for the share > private contrast in the right nucleus accumbens (NAc). (B) Negative association between PAiS score and listener-related neural modulation for the share > private contrast in the left frontal operculum (FOP4), inferior frontal gyrus pars orbitalis (IFG), and ventral posterior superior temporal sulcus (STSvp). Scatter plots display, for each adult who stutters (*n* = 33), PAiS score on the x-axis and the mean participant-level parameter estimate for the share > private contrast, averaged across all voxels within the respective significant cluster, on the y-axis. Each data point represents one participant. Analysis was carried out using a fixed-effects model, by forcing the random effect variance to zero in FLAME (FMRIB’s Local Analysis of Mixed Effects). Contrast maps are cluster-corrected using a cluster-forming threshold of *Z* > 3.1 and *p* < .05. **ALT TEXT:** Panels A and B show brain sections and scatter plots relating stuttering anticipation scores in adults who stutter to listener-related neural modulation, measured as the contrast between share and private trials. A shows a positive relationship in the right nucleus accumbens, with higher anticipation scores associated with a larger share versus private difference. B shows a negative relationship in left inferior frontal and superior temporal regions, with higher anticipation scores associated with a smaller share versus private difference.

Because trait anxiety and educational attainment can influence prefrontal and striatal engagement, we additionally conducted exploratory partial correlation analyses within the AWS group. Applying a Bonferroni-corrected alpha of 0.05/3 ≈ 0.017, stuttering anticipation remained robustly associated with striatal (*r* = 0.55, *p* < 0.001) and temporal (STS: *r* = -0.52 *p* = 0.002) modulation when controlling for trait anxiety and professional education (Supplementary Table 7). By contrast, the association with the frontal cluster (IFG BA 44, 45, 47: *r* = -0.36, *p* = 0.041) did not survive correction for multiple comparisons.

To clarify whether the PAiS-related effects in the share > private contrast reflected listener-specific modulation or different condition-specific response patterns, we extracted parameter estimates for the share and private conditions separately from the identified clusters. Correlation analyses revealed no significant condition-wise association in right NAc, a negative association with private speech only in left STS, and negative associations with both shared and private speech in left FOP4/IFG (Supplementary Fig. 7), the latter consistent with reduced recruitment across conditions and weaker listener-related differentiation at higher PAiS. Moreover, the parameter estimates of the right NAc cluster from the share > private contrast were not significantly correlated with those of the PAiS-associated left STS or left FOP4/IFG clusters (Supplementary Fig. 8), indicating that these regional effects did not covary across participants.

### Correlates with stuttering-related burden in listener-related modulation

Within AWS, greater stuttering-related burden (OASES) was positively associated with listener-related modulation of brain activity (share > private). Higher OASES scores correlated with increased responses in medial frontal and cingulate regions, including medial prefrontal cortex, anterior cingulate cortex, and medial superior frontal cortex, as well as inferior frontal and frontal opercular regions (Fig. 6; Supplementary Table 8).

**Figure 6.**
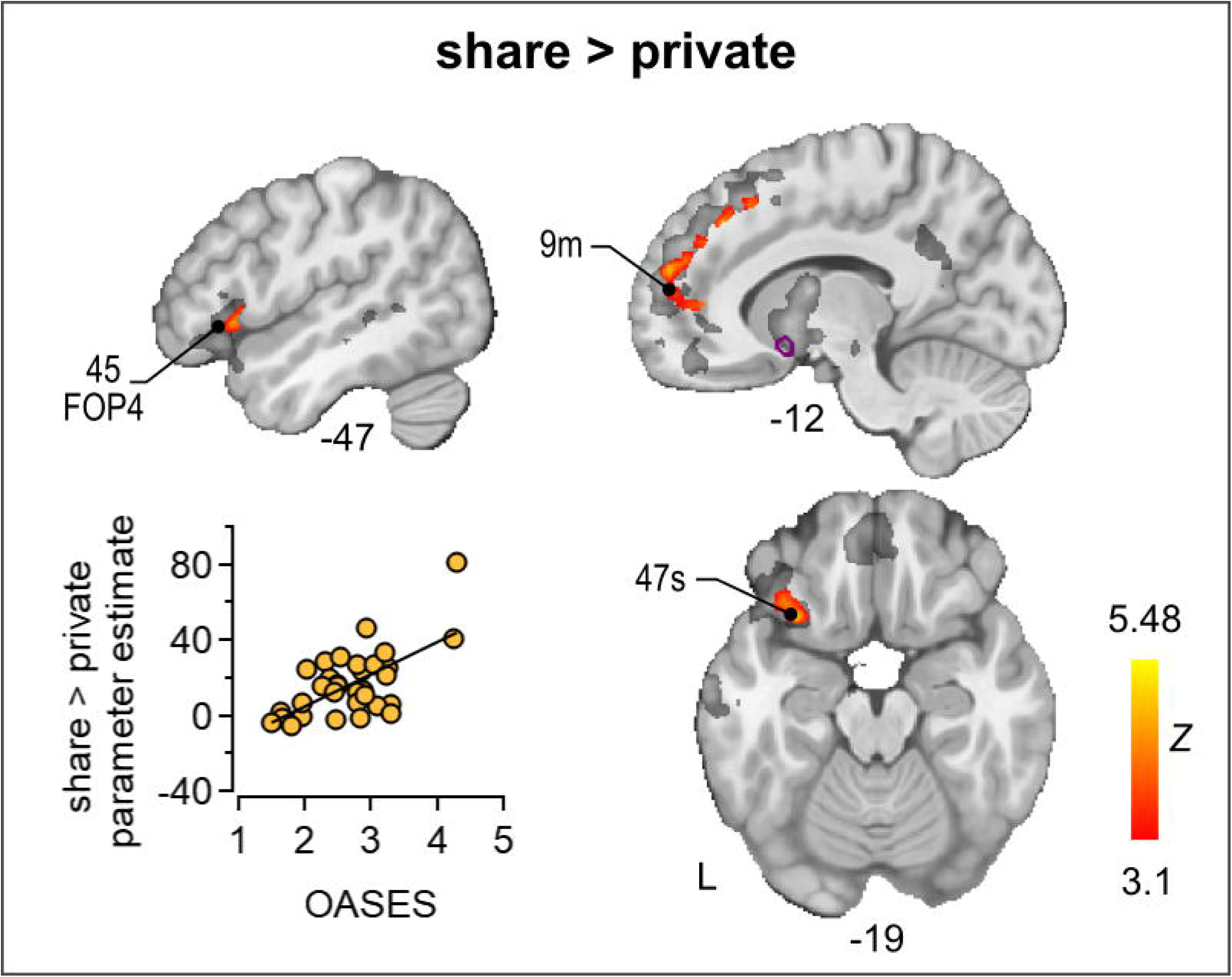
Correlates of overall stuttering experience. In adults who stutter (n = 33), within the cluster mask of the share > private contrast (dark grey overlay), higher OASES scores were positively associated with listener-related modulation in medial prefrontal and medial superior frontal cortex, anterior cingulate cortex (9m), and inferior frontal gyrus (pars orbitalis and triangularis) and frontal operculum (FOP4, 45). The scatter plot shows the relationship between OASES score and mean participant-level parameter estimates for the share > private contrast across all significant clusters; each data point represents one participant. Analysis was carried out using a fixed-effects model, by forcing the random effect variance to zero in FLAME (FMRIB’s Local Analysis of Mixed Effects). Contrast maps are cluster-corrected using a cluster-forming threshold of *Z* > 3.1 and *p* < .05. **ALT TEXT:** Brain sections and a scatter plot relating the overall impact of stuttering in adults who stutter to listener-related neural modulation, measured as the contrast between share and private trials, within regions showing a share versus private effect. Higher scores are associated with a larger share versus private difference in medial prefrontal and anterior cingulate cortex and in left inferior frontal and orbitofrontal regions.

To assess whether trait anxiety or educational attainment could explain the associations between stuttering impact and frontal cortical sites, we performed further exploratory partial correlation analyses within the AWS group. Applying a Bonferroni-corrected alpha of 0.05/6 ≈ 0.008, stuttering-related burden (OASES) remained significantly associated with left IFG (BA 47), left and right medial prefrontal cortex (left 9m/p24, right 9m), and medial superior frontal cortex (8BM/SCEF) when controlling for trait anxiety and professional education (Supplementary Table 9). By contrast, the association with left IFG pars triangularis (BA 45) showed only a trend toward significance at this threshold, and the association with left anterior cingulate cortex (d32) did not survive correction for multiple comparisons.

## Discussion

Speaking is a motor act embedded in socially motivated behavior that engages valuation and control systems. In a socio-economic decision task where participants chose between speaking and remaining silent, both listener presence and self-related speech carried clear subjective value for AWS and AWNS extending prior findings from fluent speakers ^29^ to AWS. Using fMRI, we further found that listener-directed and self-related speech engage reward- and evaluation-related brain regions, including ventral striatum and medial frontal cortex, in both groups. Although listener presence did not produce group differences, individual variability in neural modulation was systematically related to stuttering anticipation and overall impact, indicating that social context shapes brain activity in AWS along clinically relevant dimensions. Importantly, these associations were observed for trait-like measures of stuttering anticipation and everyday impact, rather than for condition-specific variation in overt stuttering frequency, suggesting that the neural effects are more closely related to enduring dispositional and experiential factors than to the immediate occurrence of stuttering events under the present task conditions.

Before turning to stuttering-specific heterogeneity, it is important to note that the main effects of listener presence and self-relatedness were robust across groups and broadly consistent with prior work linking socially contextualized communication to medial frontal and temporo-parietal regions, and in some paradigms to reward-related systems^51–54^. The temporo-parietal and medial prefrontal regions implicated by the self-related contrast fall within a broader set of areas commonly associated with self-referential and social-cognitive processing, including mentalizing-related functions, and partially overlap with regions of the default mode network^55^.

In the following discussion, we focus on aspects of the data that are specific to stuttering, namely, how inter-individual differences in anticipation of stuttering and overall stuttering impact relate to modulation of speech-related brain activity by social context. Moreover, the observed brain-behaviour associations were estimated using fixed-effects analyses and therefore formally apply only to the present sample, rather than supporting population-level inference. These exploratory correlations need to be interpreted as characterizing heterogeneity within the cohort of AWS, and as hypothesis-generating with respect to how clinically relevant dimensions of the stuttering phenotype may shape processing of speech in social contexts.

Prior structural and resting state studies already indicate that cortico basal ganglia and frontotemporal networks are altered in developmental stuttering, including anomalous resting state network architecture, atypical connectivity of speech motor and auditory regions, and morphology of the nucleus accumbens that differs between children and adults who stutter ^22,32,56,57^. Our results extend this work by showing how these large scale systems are differentially engaged by social context and stuttering load during active speech. Together with developmental MRI data showing distinct trajectories of striatal and cortical networks in children who stutter ^23^, our findings support the view that social communicative experience contributes to experience dependent reorganization of cortico basal ganglia loops over time.

### Behavioral effects of listener presence

Despite reports that AWS avoid challenging communicative situations, we did not observe systematic avoidance in this low-stakes setting: AWS did not forgo monetary reward to avoid speaking, and stuttering frequency did not differ reliably between shared and private trials. This suggest that, under these conditions, the subjective value of speaking and self disclosure remains high in AWS and can be dissociated from overt avoidance behavior. It also implies that the neural effects we observed are unlikely to be driven simply by increased dysfluency under social exposure, but instead reflect changes in the motivational and regulatory state in which speech is produced. However, the paradigm manipulated listener presence without reciprocal interaction or explicit social evaluation, and the absence of any reply from the experimenter likely limited the social relevance of the manipulation.

### Premonitory awareness, motivational valuation and speech-language hubs

Within AWS, listener-related modulation of the nucleus accumbens scaled with individual differences in stuttering anticipation: higher PAiS scores were associated with stronger differentiation between directed and private speech in this ventral striatal region. Notably, this association remained significant after controlling for trait anxiety and professional education, suggesting that it is unlikely to be explained solely by general anxious disposition or educational background. Given the established role of the nucleus accumbens in encoding motivational relevance and subjective value in socially meaningful contexts ^58–64^, this pattern suggest that socially directed speech acquires greater motivational salience as anticipatory load increases, rather than simply becoming more rewarding. In multi-system terms, anticipatory load appears to amplify automatic motivational signals in ventral striatum, consistent with how affective cues bias approach-avoid decisions via striatal circuitry ^33^. Structural work has identified altered morphology of the nucleus accumbens in persistent developmental stuttering, with reduced volume in children but increased right lateralized volume in adults ^22,32^, suggesting a non linear developmental trajectory of ventral striatal circuitry. Our finding that listener related modulation in this region scales with anticipatory load is consistent with the idea that atypical ventral striatal development is functionally expressed as heightened motivational salience of socially directed speech.

Higher stuttering anticipation was also associated with reduced listener-related modulation in cortical speech-language regions, including inferior frontal and posterior temporal areas that support speech planning and perception ^65–67^, as well as the lateral orbitofrontal cortex (BA 47), implicated in controlled semantic processing and in integration of value and context into language and decision-making ^68,69^. Follow-up analyses showed that these effects were not uniform across regions. In left STS, the negative association was driven primarily by the private condition, suggesting that the contrast effect reflected variation in private-speech recruitment. In left FOP4/IFG, by contrast, PAiS was negatively correlated with both shared and private speech, indicating a more general reduction in recruitment with increasing anticipation and a weaker listener effect at higher anticipation. Speculatively, this pattern may reflect a shift away from cortical speech-language processing toward motivational-salience systems under higher anticipatory load. Notably, parameter estimates from the right NAc cluster did not significantly correlate with those from the left STS or left FOP4/IFG clusters, suggesting that the striatal and left-hemisphere effects may reflect partially dissociable aspects of anticipation-related modulation rather than a single common process.

Recent work indicates that the core language network, although not itself computing others’ mental states ^70,71^, can be rapidly informed by feedback from theory-of-mind regions during naturalistic comprehension, with reliable functional coupling between these networks over time ^72^. Within broader prefrontal circuitry, lateral orbitofrontal and frontopolar regions have been linked to credit assignment and to counteracting biased value signals ^33,73,74^ complementing striatal mechanisms of value updating ^75^. In this context, concurrently reduced modulation in ventrolateral frontal regions, together with stronger listener-related responses in nucleus accumbens, is consistent with a shift in the relative engagement of ventral striatal motivational circuitry and cortical speech language systems as stuttering anticipation increases.

### Stuttering-related burden is associated with self-referential and regulatory regions

Listener-related increases in activity for share > private speech were observed in a medial prefrontal-cingulate network associated with social evaluation, self-referential processing, and context-dependent regulation ^76–80^. Their greater engagement for directed relative to private speech suggests that speaking under observation places additional demands on evaluative and monitoring processes. Within AWS, higher OASES scores were associated with stronger listener-related modulation in medial prefrontal and ventrolateral frontal regions, indicating that individuals who experience greater stuttering-related burden recruit this evaluative-regulatory circuitry more strongly under listener presence. Follow-up partial correlations controlling for trait anxiety and professional education showed that the main ventral striatal and ventrolateral/orbitofrontal associations remained significant, although some weaker frontal effects did not survive correction for multiple comparisons. Because OASES and trait anxiety were strongly correlated, these analyses should be interpreted conservatively: they support a relation to stuttering-related anticipatory and evaluative burden beyond general anxiety, but do not fully dissociate motivational valuation, cognitive effort, and emotional distress. Notably, this ventrolateral/orbitofrontal involvement also resonates with prior work implicating left BA 47 in recovery from developmental stuttering.^81,82^ In recovered speakers, orbitofrontal and superior cerebellar regions were reported to become functionally uncoupled from the broader speech production network.^81^ Taken together, these findings suggest that orbitofrontal cortex is more sensitive to social context in individuals who experience stuttering as more burdensome in everyday life, and raise the possibility that this region helps mediate social-context effects on speech production. In this view, recovery may involve neuroplastic reorganization of orbitofrontal cortex and its interaction with speech production circuitry, including superior cerebellar regions, although this interpretation remains tentative.

OASES and PAiS were not correlated (Spearman *r*(31) = 0.27, *P* = 0.13, two-tailed), supporting the view that these neural associations track distinct aspects of the stuttering experience and phenotype. Within a multi-system framework ^33^, dorsal anterior cingulate cortex and medial prefrontal cortex may act as a hub, where learned negative consequences of speaking and current contextual cues jointly shape the value of speaking under observation.

The self > other contrast revealed group differences in regions implicated in self-referential and evaluative processing. Relative to AWS, AWNS showed greater engagement of the subcallosal cortex, often linked to affective valuation and self-related appraisal ^83,84^, whereas AWS exhibited stronger activity in the medial inferior parietal cortex (PFm), a region near the temporo-parietal junction that lies within a broader parietal territory commonly implicated in self-referential processing, perspective-taking, integration of self-related information within a broader social context, ^85,86^ and partial default mode network overlap. More generally, the self > other main effect engaged medial prefrontal and parietal regions consistent with prior work on self-related thought and social cognition^55^. Although these differences require replication, the pattern suggests that self-related speech may rely on partially distinct self-referential circuits in AWS and AWNS, potentially reflecting differences in how self-relevant information is evaluated or contextualized during speaking. On current accounts, ventromedial prefrontal cortex/subcallosal regions encode cognitive ‘maps’ for self relevant value, whereas parietal regions support perspective taking and contextual integration ^33^. Within this view, self related speech in AWS may depend relatively more on distributed perspective taking networks and relatively less on ventromedial prefrontal centered self valuation schemas, possibly reflecting experience dependent adaptations to chronic social evaluative contexts. However, the parietal group difference should be interpreted cautiously, as it was attenuated after adjustment for trait anxiety and professional education. Together with the OASES correlations, these group differences indicate that both the evaluation of self-related speech and its modulation by listener presence vary systematically with the perceived impact of stuttering.

This dovetails with evidence that, after fluency-shaping therapy, greater relief from the social-emotional burden of stuttering is linked to lower fractional anisotropy in the right frontal aslant tract, suggesting that both frontal grey and white matter systems contribute to how stuttering burden is experienced and reduced. Consistent reports of altered frontotemporal and cerebellar-orbitofrontal connectivity that scale with stuttering severity ^56^ further suggest that individuals with higher burden recruit evaluative and regulatory hubs within a broader fronto-cerebellar and temporo parietal network. In our task, this is reflected in stronger context dependent engagement of medial prefrontal and ventrolateral frontal regions when speaking under observation.

Prior work using naturalistic interpersonal communication paradigms has shown that disfluent speech in adults who stutter is accompanied by right amygdala activity and reduced prefrontal engagement ^18^, highlighting a prominent role for limbic circuits during socially demanding speech. Our results complement these findings by showing that, even in a relatively low stakes setting, higher stuttering-related burden shifts listener related modulation toward medial prefrontal und lateral orbitofrontal valuation circuits. Together, these studies indicate that limbic, cognitive and sensorimotor systems jointly shape how social context is mapped onto brain networks in developmental stuttering.

Overall, our findings do not challenge the contribution of sensorimotor abnormalities to developmental stuttering, but instead suggest that these abnormalities are embedded within a broader, context-sensitive network in which salience, control, and language processing systems are differentially engaged as a function of anticipation and burden. This system level perspective helps explain why stuttering is highly context sensitive and heterogeneous and suggests that targeting motivational and evaluative processes, alongside speech motor mechanisms, may be important for understanding and treating the disorder.

### Limitations

A methodological consideration concerns the ecological validity of the PSE as a measure of communicative burden. The monetary trade-offs employed here, following Tamir & Mitchell (2012), are deliberately small and serve as a utility metric to reveal preference structure through real choices, rather than to represent the absolute psychosocial cost of stuttering in social life. For many individuals who stutter, the stakes of speaking in social contexts, including stigma, anticipatory anxiety, and communicative failure, far exceed what can be offset by small financial incentives in a laboratory setting. To the extent that adults who stutter experienced the task as lower-stakes than naturalistic communication, the observed group differences in PSE and in neural modulation may represent artificial estimates of real-world disparities. This does not invalidate the within-paradigm comparisons, which remain internally consistent, but caution is warranted when extrapolating PSE values to real-world communicative costs.

The paradigm isolates listener presence and self-relatedness but does not directly manipulate social evaluation or communicative consequences, and the reported associations are correlational rather than causal. Furthermore, the private condition may not constitute a fully socially neutral baseline, as residual awareness of the experimental context could maintain some evaluative load for individuals who stutter. Conversely, the supportive context of participating in a stuttering study may have attenuated the evaluative load of share trials relative to naturalistic listener-directed speech. Both conditions may thus represent intermediate points on a social evaluation continuum, and the contrast between them should be interpreted as a measure of relative rather than absolute social modulation. More naturalistic interaction paradigms and longitudinal designs will be needed to clarify how motivational and regulatory systems interact with speech-motor processes over time ^87^. Correlation analyses were conducted using a fixed effects model in FSL, confining inference to the present sample. This choice was motivated by the heterogeneity of the stuttering phenotype, high inter-individual variability in neural responses, and the absence of significant random effects for the critical contrast. Generalizability to the broader population therefore awaits replication in larger, independent cohorts.

## Conclusion

Overall, our findings do not argue against sensorimotor accounts of developmental stuttering, but show that communicative context and clinically relevant individual differences additionally shape speech-related brain activity with stronger links to trait-like measures of anticipation and everyday burden than to overt stuttering frequency under the present task conditions. This multi-system perspective may help account for the context sensitivity and heterogeneity of stuttering and underscores the contribution of social and emotional factors to the neural organization of speech processing. Our findings do not directly speak to treatment efficacy but suggest that clinical assessment may benefit from explicitly considering how social exposure and stuttering anticipation modulate activity in salience, control, and speech-language systems, alongside traditional measures of overt fluency. This may help explain why the same individual can show very different fluency profiles across situations and support approaches that focus on reducing social and emotional load in challenging contexts rather than targeting speech motor behavior in isolation.

## Supplementary material

Supplementary material is available at *Brain Communications* online.

## Data availability

Data availability: Custom code for the processing of the multi echo fMRI data is available at: https://doi.org/10.25625/LZJNSZ. The brain imaging data of the group averages in the standard space are available at: https://doi.org/10.25625/OQBVSL.

## Acknowledgements

We are grateful for the generous support of Angela D. Friederici. Additionally, we wish to thank all of our participants for volunteering their time.

## CRediT statements

Nicole E. Neef: Conceptualization, Investigation, Data curation, Formal analysis, Project administration, Visualization, Supervision, Validation, Writing - original draft, Writing - review & editing. Elena Winter: Formal analysis, Writing - review & editing. Andreas Neef: Methodology, Formal analysis, Writing - review & editing. Toralf Mildner: Methodology, Writing - review & editing. Shaza Haj Mohamad: Formal analysis, Writing - review & editing. Hellmuth Obrig: Investigation, Writing - review & editing. Christian H. Riedel: Resources, Writing - review & editing. Katharina Scholze: Investigation, Data curation, Formal analysis, Writing - review & editing.

