## Supplementary Figures for "Striatal and frontal signatures of social context and cost-benefit decision making in developmental stuttering"

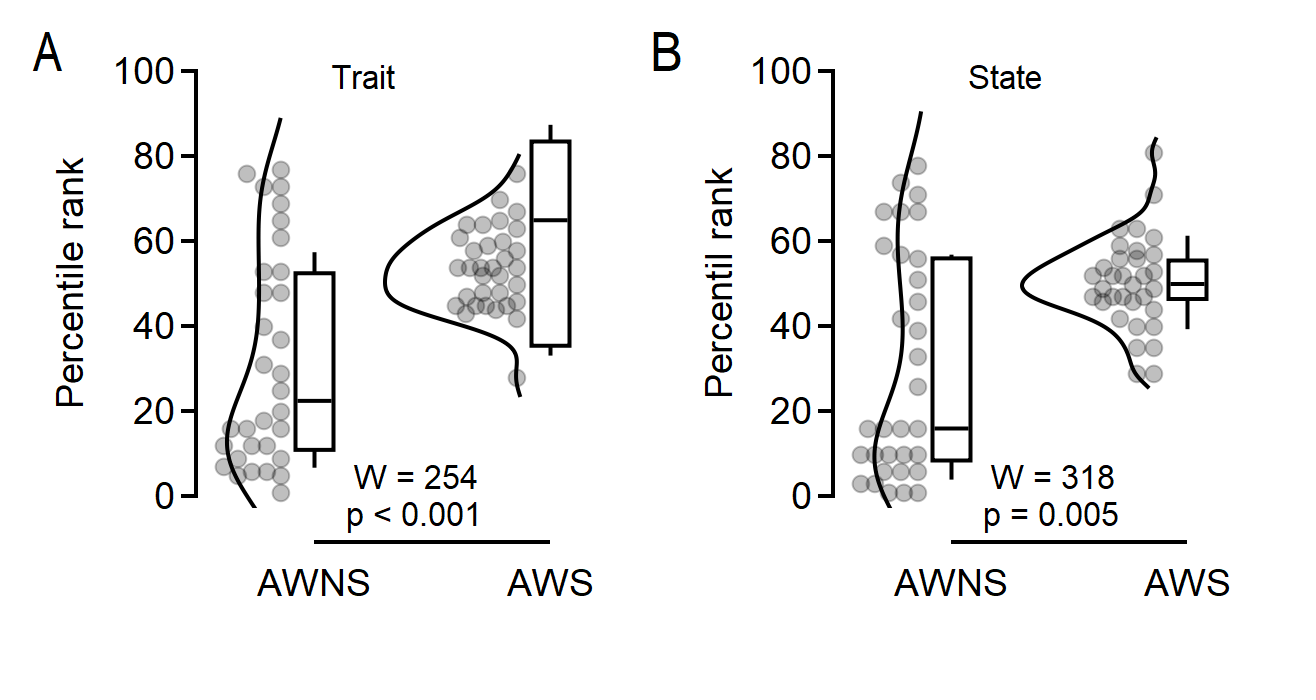

**Supplementary Figure 1.** (A) Distribution of trait-anxiety percentile ranks in adults who do not stutter (AWNS, *n* = 32) and adults who stutter (AWS, *n* = 33). (B) Distribution of state-anxiety percentile ranks in AWNS and AWS. Each grey dot represents the percentile-rank score of one participant. Half-violin plots illustrate the distributions; box plots show the median (central horizontal line) and interquartile range (box), with whiskers extending to the minimum and maximum values. Group differences were tested using two-sided Wilcoxon rank-sum tests. AWS showed higher trait-anxiety percentile ranks than AWNS and higher state-anxiety percentile ranks than AWNS.

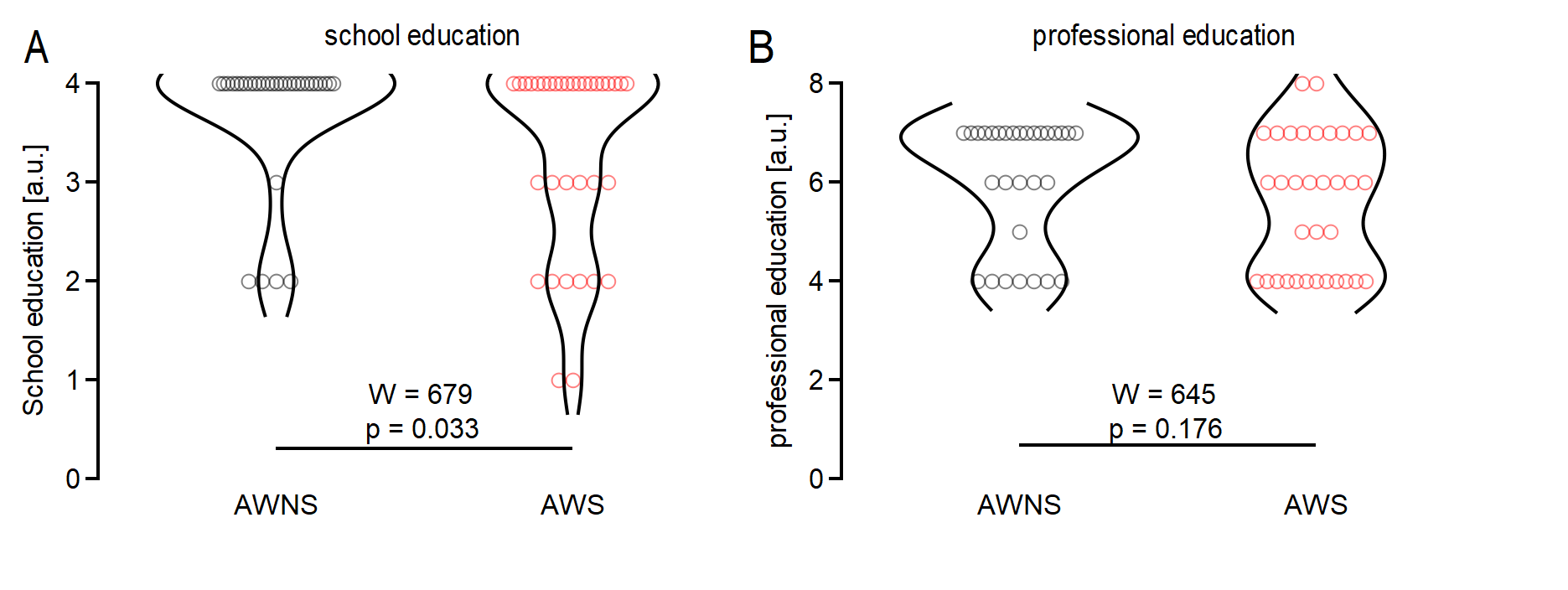

**Supplementary Figure 2.** Violin plots showing the distribution of school (A) and professional (B) education scores for adults who do not stutter AWNS (grey, *n* = 32) and adults who stutter AWS (red, *n* = 34) in the samples. School education scores are markedly skewed, with most participants in both groups at the highest level. Wilcoxon rank‑sum tests indicate a small group difference in school education and no group difference in professional education. Each circle represents the education score of one participant. Highest school qualification was classified as: (0) no qualification, (1) general education school-leaving certificate after grade 9, (2) general education school-leaving certificate after grade 10, (3) university of applied sciences entrance qualification, and (4) higher education entrance qualification. Highest professional qualification was classified as: (0) no qualification; (1) vocational training <1 year; (2) vocational training <2 years; (3) vocational training <3 years; (4) completed vocational qualification; (5) currently enrolled at a university or university of applied sciences; (6) bachelor’s degree or master craftsman qualification; (7) master’s degree or equivalent; and (8) doctoral degree or higher.

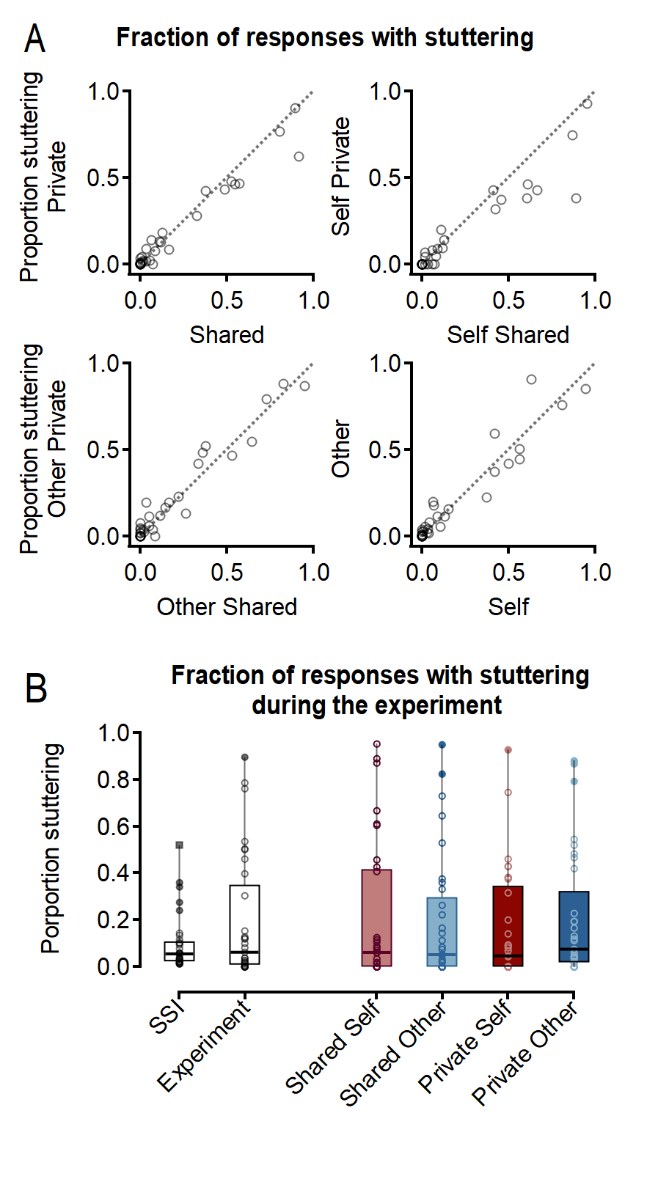

**Supplementary Figure 3. Fraction of responses with stuttering.** (A) Scatterplots compare the fraction of responses with stuttering between the shared and private conditions (overall and separately for self and other) and between the self and other conditions in adults who stutter (*n* = 34). Each circle represents one participant. The dotted line indicates the identity line (x = y); points below the line indicate more stuttering in the condition on the x-axis, and points above the line more stuttering in the condition on the y-axis. The scatter plots are shown for illustration only; no correlation analysis was performed. In the experiment, private speech did not significantly reduce the occurrence of stuttering (Wilcoxon signed-rank tests; *self share* and *self private*: *V* = 308, *p* = .051; *other share* and *other private*: *V* = 200, *p* = .155, uncorrected; see Supplementary Table 2). (B) Distribution of participant-level proportions of responses containing stuttering during the SSI-4 assessment, during the experiment collapsed across conditions, and separately for the shared-self, shared-other, private-self, and private-other conditions. Each circle represents the proportion of responses containing stuttering for one AWS in the respective assessment or experimental condition. Boxes indicate the interquartile range, horizontal lines indicate the median, and whiskers indicate the minimum and maximum values.

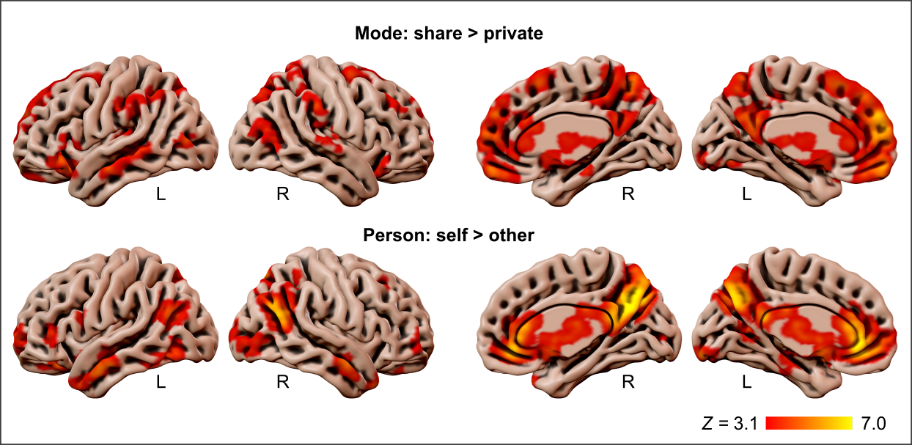

**Supplementary Figure 4. Surface renderings of main effects across all participants** (*n* = 66). Lateral and medial views of the left and right hemispheres showing activation for the share > private (top) and self > other (bottom) contrasts across all participants. All fMRI results are based on FLAME (FMRIB’s Local Analysis of Mixed Effects) stage 1. Contrast maps are cluster-corrected using a cluster-forming threshold of Z > 3.1 and p < .05. Peak coordinates and cluster statistics are reported in Supplementary Table 3.

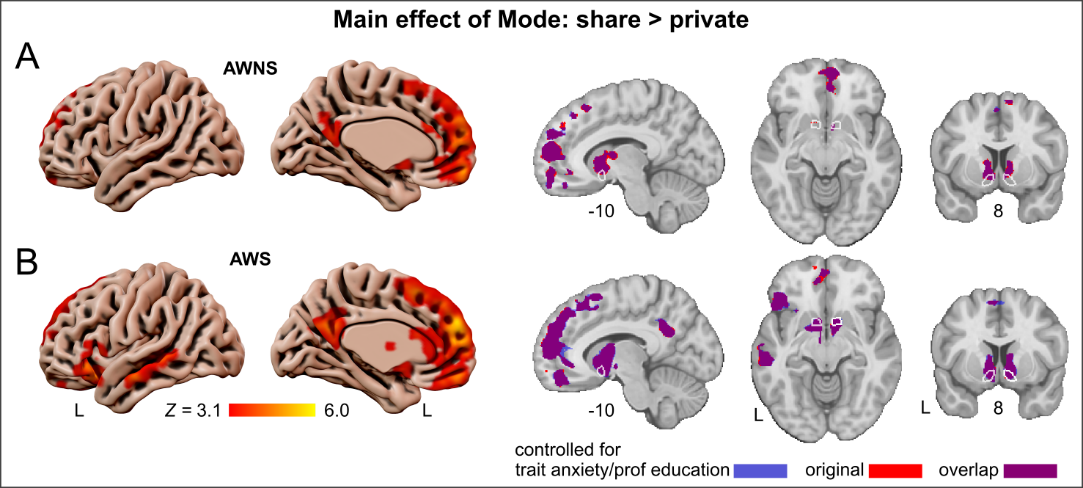

**Supplementary Figure 5. Listener presence controlled for trait anxiety and professional education.** Whole-brain results for the share > private contrast, separately for (A) adults who do not stutter (AWNS, *n* = 32) and (B) adults who stutter (AWS, *n* = 33), after including trait anxiety (STAI) percentile rank and professional education as demeaned covariates of no interest in the group-level model. Surface renderings (left panels) display cluster extent with Z-statistic color bars; brain slices (right panels) highlight voxels significant only in the covariate model (blue), only in the original model (red), or in both (purple). All fMRI results are based on FLAME (FMRIB’s Local Analysis of Mixed Effects) stage 1. All maps are thresholded using cluster-wise correction (cluster-forming threshold Z > 3.1, cluster-extent p < 0.05, family-wise error corrected). The spatial pattern and extent of activations are consistent with the uncontrolled results shown in Figure 3, indicating that listener-related modulation of the nucleus accumbens, caudate nucleus, and anterior cingulate cortex is not attributable to between-group differences in trait anxiety or professional education level. An exception was observed in AWNS, where thalamic listener-related modulation was reduced after covariate adjustment, suggesting that thalamic responses to listener presence are partially explained by individual differences in trait anxiety or professional education.

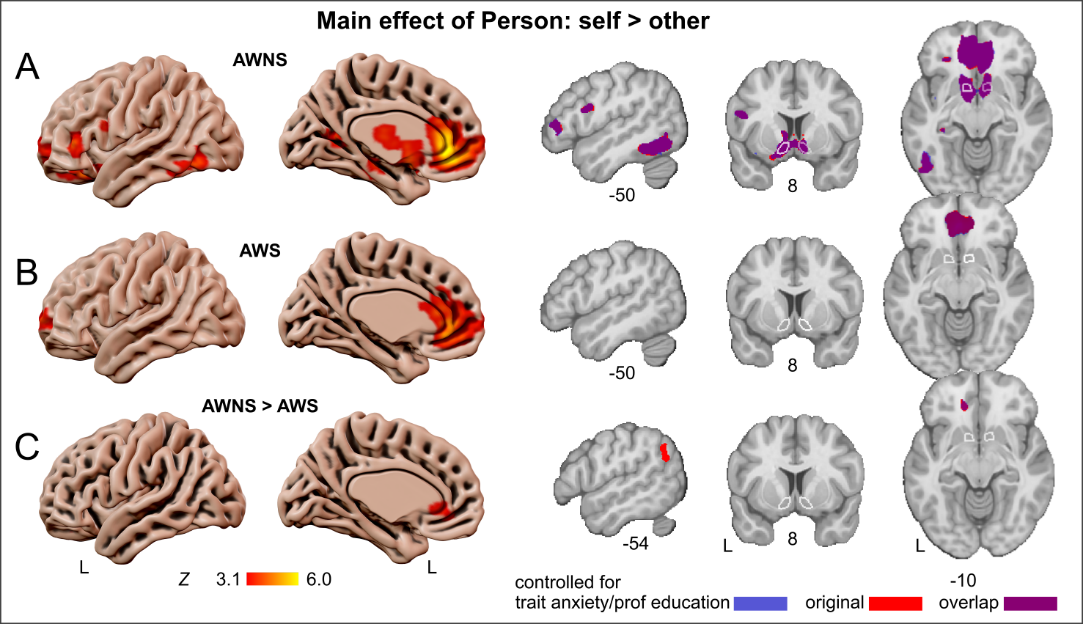

**Supplementary Figure 6. Self-related speech controlled for trait anxiety and professional education.** Whole-brain results for the self > other contrast, separately for (A) adults who do not stutter (AWNS, *n* = 32) and (B) adults who stutter (AWS, *n* = 33), after including trait anxiety (STAI) percentile rank and professional education as demeaned covariates of no interest in the group-level model. Surface renderings (left panels) display cluster extent with Z-statistic color bars; brain slices (right panels) highlight voxels significant only in the covariate model (blue), only in the original model (red), or in both (purple). All fMRI results are based on FLAME (FMRIB’s Local Analysis of Mixed Effects) stage 1. All maps are thresholded using cluster-wise correction (cluster-forming threshold Z > 3.1, cluster-extent p < 0.05, family-wise error corrected). The spatial pattern and extent of activations are consistent with the uncontrolled results shown in Figure 4, indicating that person-related modulation of the nucleus accumbens, caudate nucleus, and anterior cingulate cortex is not attributable to between-group differences in trait anxiety or professional education level. An exception was observed in the group contrast, where the reduced activation of the left inferior parietal cortex (area PFm) in AWS did not survive covariate adjustment.

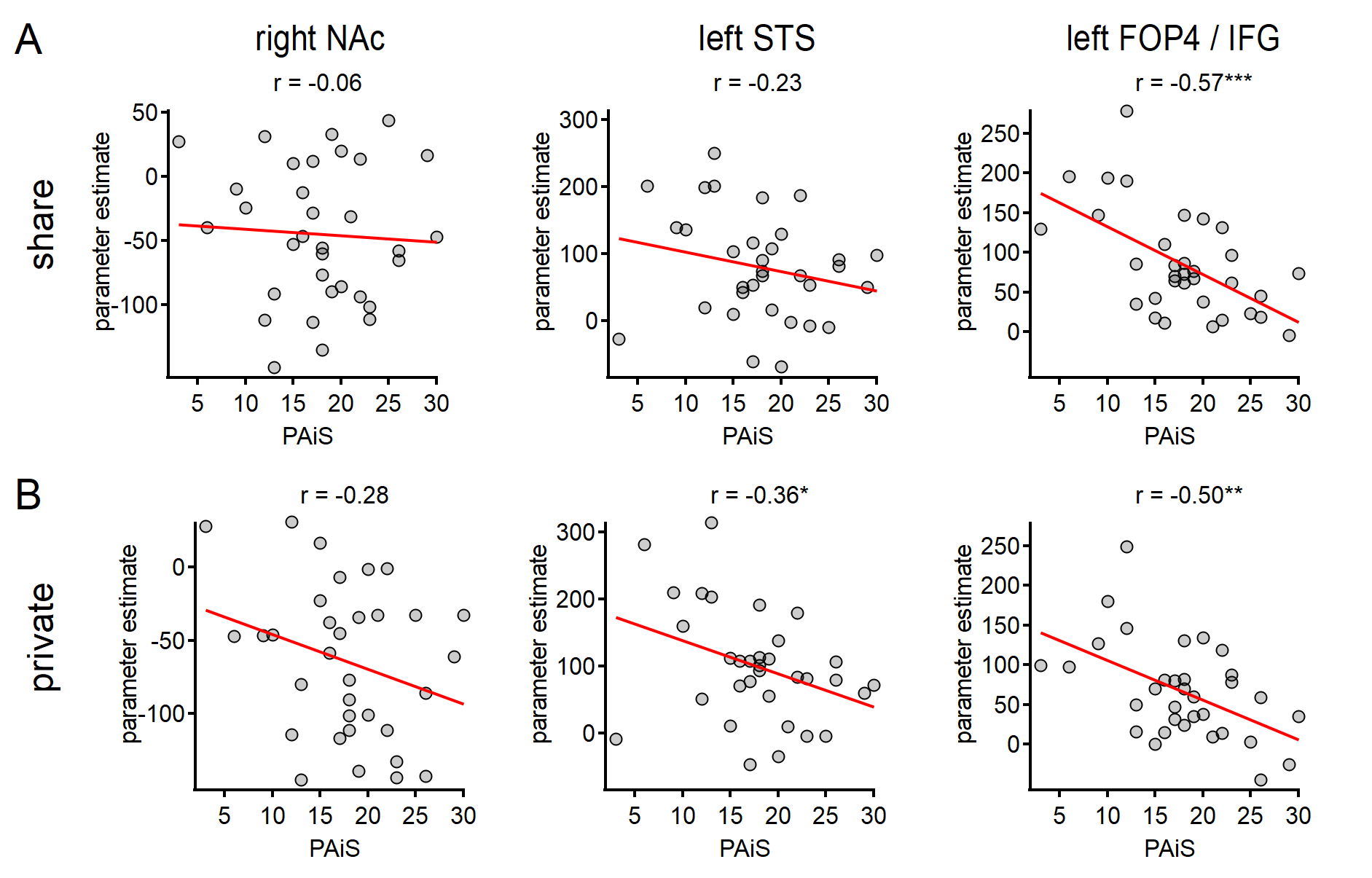

**Supplementary Figure 7. Exploration of the associations between PAiS and parameter estimates separated for share and private conditions in the striatal and speech clusters.** Scatter plots show linear associations between PAiS score and parameter estimates extracted separately for the shared (A) and private (B) speaking conditions in adults who stutter (AWS; 𝑛 = 33). Columns show the right nucleus accumbens (NAc), left superior temporal sulcus (STS), and left frontal operculum/inferior frontal gyrus (FOP4/IFG) clusters, respectively. Each circle represents one AWS. The x-axis shows the participant’s PAiS score, and the y-axis shows that participant’s mean parameter estimate for the displayed condition, averaged across voxels within the respective cluster. Red lines show linear regression fits. Associations were assessed using two-sided Pearson product–moment correlations; the displayed values are Pearson’s 𝑟; significance is indicated as follows: **p* < 0.05, ***p* < 0.01, ****p* < 0.001.

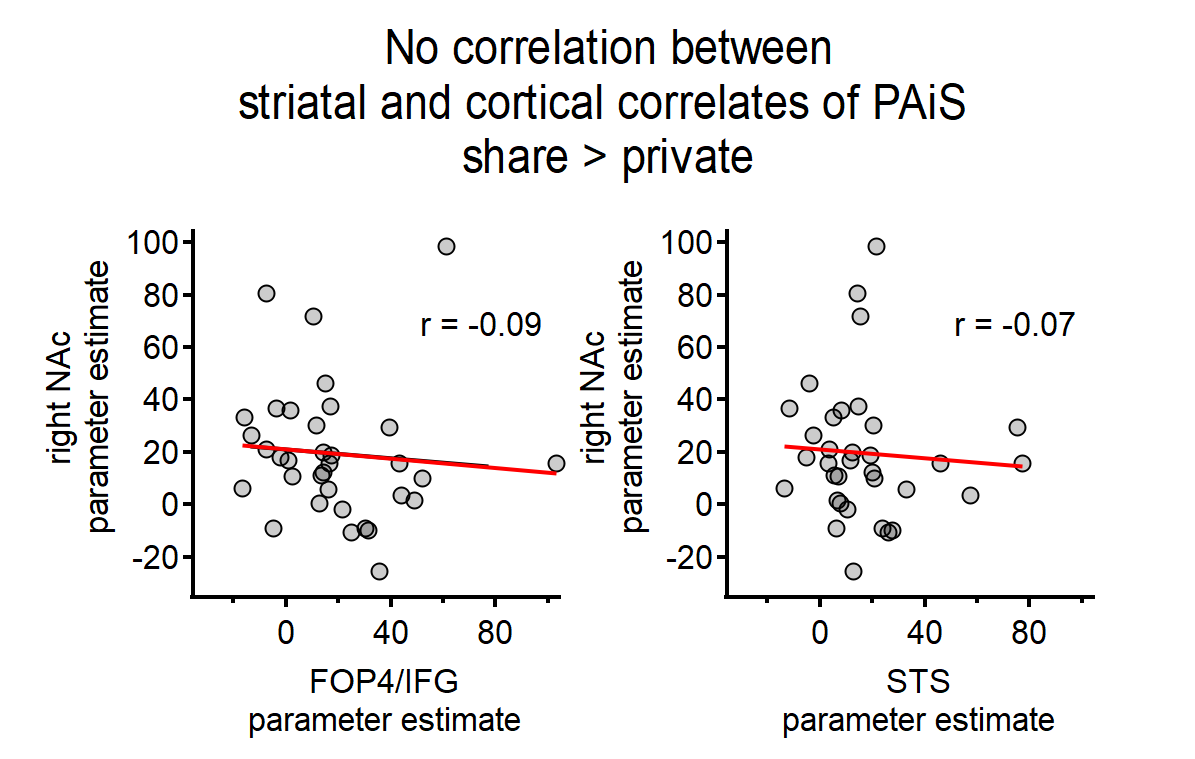

**Supplementary Figure 8. Exploration of the association between striatal and cortical contrast parameter estimates in the clusters significantly correlated with PAiS.** Scatter plots show the relationship between participant-level parameter estimates for the share > private contrast in the right nucleus accumbens (NAc) and those in the left frontal operculum/inferior frontal gyrus (FOP4/IFG) and left superior temporal sulcus (STS) in adults who stutter (AWS; 𝑛 = 33). Each circle represents one AWS. The x-axis shows the participant’s mean parameter estimate for the share > private contrast, averaged across voxels within the respective cortical cluster; the y-axis shows the corresponding mean parameter estimate averaged across voxels within the right NAc cluster. Red lines indicate linear regression fits. Associations were assessed using two-sided Pearson product-moment correlations. Neither association was statistically significant (*p* > .05).
