## Supplementary material for "Striatal and frontal signatures of social context and cost-benefit decision making in developmental stuttering"

**Tables**

**Supplementary Table 1** Relative monetary value for each condition

|  | Mean [95% CI]  Fluent | Mean [95% CI]  Stuttering | *T* | *p_uncor_* | *p_cor_* |
| --- | --- | --- | --- | --- | --- |
| Self Shared | 0.98 [0.53, 1.44] | 1.76 [1.27, 2.25] | -2.38 | .021 | .083 |
| Other Shared | 1.00 [0.58, 1.42] | 1.59 [1.09, 2.09] | -1.84 | .071 | .283 |
| Self Private | 0.77 [0.41, 1.14] | 1.31 [0.84, 1.78] | -1.83 | .072 | .290 |
| Other Private | 0.57 [0.22, 0.93] | 1.21 [0.76, 1.66] | -2.29 | .026 | .103 |

^Mean and 95 % confidence interval for each condition and group. Differences tested statistically with unpaired t-tests (R command: t.test) uncorrected and with Bonferroni correction.^

**Supplementary Table 2** Fraction of stuttered responses in percent

|  |  | Median (IQR)  Shared | Median (IQR)  Private | *V* | *p_uncor_* | *p_cor_* |
| --- | --- | --- | --- | --- | --- | --- |
| Stuttering | Self | 6.12% (41.73%) | 4.65% (34.66%) | 308 | .051 | .101 |
|  | Other | 5.17% (29.76%) | 7.50% (30.52%) | 200 | .155 | .309 |
| Control | Self | 0% (0%) | 0% (0.52%) | 12 | .461 | .922 |
|  | Other | 0% (0.48%) | 0% (2.22%) | 18 | 1 | 1 |

^Differences tested statistically with paired Wilcoxon Signed-Rank Tests (R command: wilcox.exact) uncorrected and with Bonferroni correction.^

**Supplementary Table 3** Whole-brain *F*-test results for Mode (share > private) and Person (self > other) effects (*Z* ≥ 3.1; *P* = 0.05, cluster-corrected, grey matter masked).

| **Region** |  | **HCPex** | **x** | **y** | **z** | ***Z*** | **Cluster size** |
| --- | --- | --- | --- | --- | --- | --- | --- |
| *Effect of Mode: share > private* |  |  |  |  |  |  |  |
| **Frontal** |  |  |  |  |  |  |  |
| Dorsomedial prefrontal cortex | L | 9m | -8 | 56 | 17 | 6.97 | 37272 |
| - Medial prefrontal cortex | L | 8BM | -3 | 19 | 54 | 5.69 |  |
| - Frontal pole | L | 10v | -2 | 38 | -30 | 3.69 |  |
| Inferior frontal cortex | L | 45* | -42 | 28 | -11 | 5.16 | 6976 |
| - Frontal orbital cortex | L | 47s | -44 | 22 | -19 | 4.96 |  |
| - Frontal orbital cortex | L | 47l | -43 | 28 | -4 | 4.85 |  |
| Ventrolateral prefrontal cortex | R | 47s | 30 | 18 | -20 | 4.51 | 1229 |
| Dorsolateral prefrontal cortex | R | 9-46d | 34 | 49 | 32 | 5.07 | 1174 |
| - Dorsolateral prefrontal cortex | R | 46 | 36 | 46 | 33 | 4.90 |  |
| - Dorsolateral prefrontal cortex | R | 8Ad | 30 | 31 | 47 | 4.52 |  |
| Premotor cortex | R | 6a | 26 | -1 | 64 | 4.91 | 939 |
| - Superior frontal gyrus | R | 6ma | 27 | 10 | 60 | 4.54 |  |
| - Dorsolateral prefrontal cortex | R | i6-8 | 28 | 2 | 62 | 4.28 |  |
| Superior frontal gyrus | L | 6a | -23 | 7 | 57 | 4.44 | 753 |
| - Superior frontal gyrus | L | 6ma | -22 | 5 | 61 | 4.07 |  |
| **Parietal** |  |  |  |  |  |  |  |
| Posteromedial parietal cortex | R | 7Am | 10 | -65 | 60 | 5.33 | 8128 |
| - Posterior parietal cortex | R | 7Pm | 8 | -63 | 53 | 5.14 |  |
| Posterior cingulate cortex | L | d23ab | -3 | -53 | 24 | 5.79 | 6127 |
| - Posterior cingulate cortex | R | d23ab | 2 | -52 | 28 | 5.73 |  |
| - Parieto-occipital sulcus | L | POS1 | -4 | -53 | 10 | 4.66 |  |
| Angular gyrus | L | PFm* | -47 | -52 | 44 | 5.41 | 2984 |
| - Intraparietal sulcus | L | IP2 | -45 | -48 | 44 | 5.09 |  |
| - Anterior intraparietal sulcus | L | AIP | -36 | -46 | 34 | 4.75 |  |
| Superior parietal lobule | L | 7A* | -13 | -65 | 51 | 5.19 | 3116 |
| - Superior parietal lobule | L | 7Am | -11 | -63 | 63 | 5.15 |  |
| - Posterior parietal cortex | L | 7Pm | -6 | -67 | 52 | 4.71 |  |
| Posterior cingulate gyrus | R | 5Ci | 12 | -37 | 40 | 5.30 | 2187 |
| - Posterior cingulate cortex | R | 31a | 4 | -39 | 45 | 3.88 |  |
| Supramarginal gyrus | L | PF* | -56 | -34 | 35 | 4.35 | 1660 |
| - Anterior supramarginal gyrus | L | PFt | -60 | -21 | 39 | 4.29 |  |
| - Supramarginal gyrus | L | PF | -59 | -37 | 39 | 3.99 |  |
| Anterior supramarginal gyrus | R | PF | 64 | -33 | 37 | 5.01 | 1275 |
| Superior parietal lobule | R | 5L | 19 | -47 | 72 | 4.17 | 929 |
| - Superior parietal lobule | R | 7PC | 33 | -44 | 63 | 3.61 |  |
| - Posterior bank of the postcentral gyrus | R | 2 | 22 | -41 | 63 | 3.59 |  |
| - Superior parietal lobule | R | 7AL | 27 | -51 | 63 | 3.55 |  |
| Posterior cingulate gyrus | L | 23c | -11 | -34 | 43 | 4.24 | 663 |
| Primary somatosensory cortex | R | 2* | 32 | -41 | 49 | 4.03 | 749 |
| - Anterior intraparietal sulcus | R | AIP | 35 | -37 | 42 | 3.78 |  |
| - Intraparietal sulcus | R | IP2 | 49 | -34 | 52 | 3.54 |  |
| **Temporal** |  |  |  |  |  |  |  |
| Middle temporal gyrus | L | TE1a | -69 | -8 | -17 | 5.06 | 4252 |
| - Middle temporal cortex | L | TE1a | -66 | -12 | -17 | 4.92 |  |
| - Superior temporal sulcus | L | STSvp | -59 | -34 | -4 | 4.77 |  |
| - Middle temporal cortex |  | TE1p | -65 | -40 | 2 | 4.57 |  |
| Superior temporal gyrus | R | PBelt | 52 | -30 | 17 | 4.30 | 1485 |
| - Retroinsular cortex | R | RI | 56 | -30 | 17 | 3.87 |  |
| - Superior temporal gyrus | R | A4 | 65 | -24 | 9 | 3.85 |  |
| - Parietal operculum | R | PFcm | 48 | -30 | 20 | 3.85 |  |
| Superior temporal / parainsular  cortex | L | 52 | -37 | -23 | 2 | 4.57 | 782 |
| - Posterior insula | L | PoI1 | -37 | -21 | 3 | 4.55 |  |
| - Medial Belt Complex | L | MBelt | -35 | -25 | 9 | 4.24 |  |
| - Heschl’s gyrus | L | A1 | -41 | -28 | 15 | 3.55 |  |
| **Occipital** |  |  |  |  |  |  |  |
| Occipital fusiform gyrus | L | V8 | -26 | -77 | -10 | 5.70 | 2910 |
| - Occipital fusiform gyrus | L | V4 | -21 | -84 | -8 | 5.43 |  |
| - Occipital pole | L | V1 | -7 | -102 | -9 | 3.5 |  |
| Lateral occipital cortex | R | MST | 43 | -68 | 8 | 4.22 | 2027 |
| - Lateral occipital cortex | R | LO3 | 42 | -73 | 16 | 4.00 |  |
| Lateral occipitotemporal cortex | L | V4t | -52 | -80 | 9 | 5.33 | 820 |
| **Subcortical** |  |  |  |  |  |  |  |
| Caudate nucleus | L | Caud | -8 | 10 | -4 | 6.82 | 14010 |
| - Caudate nucleus | R | Caud | 8 | 12 | -3 | 6.42 |  |
| Cerebellum Crus I | R |  | 21 | -83 | -31 | 4.63 | 1363 |
| - Cerebellum Crus II | R |  | 21 | -80 | -33 | 4.57 |  |
| Cerebellum Crus I | L |  | -32 | -84 | -34 | 4.31 | 1045 |
| - Cerebellum Crus II | L |  | -31 | -84 | -38 | 4.11 |  |
| *Effect of Person: self > other* |  |  |  |  |  |  |  |
| **Frontal** |  |  |  |  |  |  |  |
| Anterior cingulate cortex | L | a24 | -5 | 37 | -7 | 8.31 | 38309 |
| - Anterior cingulate cortex | L | p24 | -3 | 40 | 4 | 7.84 |  |
| Anterior inferior frontal sulcus | L | IFSa | -48 | 34 | 8 | 4.25 | 859 |
| - Inferior frontal gyrus, posterior pars orbitalis | L | p47r | -49 | 40 | 5 | 4.15 |  |
| Lateral orbitofrontal cortex | L | 13l | -23 | 33 | -20 | 5.63 | 2263 |
| - Lateral orbitofrontal cortex | L | 47m | -30 | 34 | -14 | 5.16 |  |
| Ventrolateral/orbitofrontal  prefrontal cortex | R | a47r | 46 | 55 | -4 | 4.50 | 703 |
| - Dorsolateral prefrontal cortex |  | a9-46vr | 44 | 57 | 9 | 3.74 |  |
| Middle frontal gyrus | R | 8Ad | 27 | 27 | 46 | 4.57 | 667 |
| Inferior frontal gyrus | L | 44 | -52 | 12 | 21 | 4.55 | 656 |
| - ventral-rostral part of the lateral premotor cortex | L | 6r | -48 | 8 | 16 | 3.58 |  |
| Orbitofrontal cortex | R | OFC | 14 | 39 | -23 | 3.72 | 84 |
| Orbitofrontal cortex | L | OFC | -13 | 41 | -20 | 3.61 | 65 |
| **Parietal** |  |  |  |  |  |  |  |
| Precuneus cortex | R |  | 6 | -64 | 30 | 8.97 | 38833 |
| - Precuneous cortex | R | 7m | 6 | -67 | 32 | 8.83 |  |
| - Precuneous cortex | L | POS2 | -1 | -71 | 37 | 8.34 |  |
| - Precuneous cortex | R | POS2 | 6 | -69 | 37 | 8.24 |  |
| - Posterior cingulate gyrus | R | RSC | 5 | -43 | 16 | 7.48 |  |
| Angular gyrus | L | PGi | -50 | -66 | 22 | 5.83 | 6112 |
| - Temporoparieto-occipital junction | L | TPOJ2 | -56 | -68 | 22 | 5.73 |  |
| Parieto-occipital sulcus | L | POS1 | -15 | -62 | 8 | 4.87 | 1305 |
| Temporoparieto-occipital  junction | R | TPOJ3 | 52 | -63 | 23 | 7.44 | 11054 |
| - Angular gyrus | R | PGi | 51 | -67 | 31 | 6.52 |  |
| - Angular gyrus | R | PGs | 43 | -76 | 40 | 4.29 |  |
| Inferior parietal lobule | R | PFm* | 47 | -47 | 58 | 4.20 | 936 |
| - Superior parietal lobule | R | 7PC | 45 | -48 | 49 | 4.1 |  |
| - Angular gyrus | R | PFm | 47 | -50 | 51 | 3.89 |  |
| **Temporal** |  |  |  |  |  |  |  |
| Middle temporal gyrus | L | TE1a | -67 | -12 | -21 | 5.91 | 5849 |
| - Inferior temporal gyrus | L | TE2a | -57 | 10 | -35 | 4.32 |  |
| Temporal fusiform gyrus | L | TE1p | -52 | -55 | -19 | 6.15 | 4613 |
| - Lateral fusiform gyrus | L | FFC | -49 | -63 | -14 | 5.17 |  |
| - Posterior middle temporal gyrus | L | PHT | -52 | -69 | -13 | 5.04 |  |
| - Posterior inferior temporal gyrus | L | PH | -46 | -69 | -8 | 4.54 |  |
| Middle temporal gyrus | R | TE1a | 62 | -4 | -19 | 5.94 | 3350 |
| - Superior temporal sulcus | R | STSva | 55 | -7 | -17 | 5.00 |  |
| - Inferior temporal gyrus | R | TE2a | 57 | -13 | -23 | 3.41 |  |
| Dorsal temporal gyrus | R | TGd | 42 | 19 | -42 | 6.39 | 1533 |
| **Occipital** |  |  |  |  |  |  |  |
| Occipital pole | L | V2 | -11 | -95 | -12 | 6.50 | 3312 |
| - Occipital fusiform gyrus | L | V3 | -19 | -88 | -11 | 4.80 |  |
| - Lingual gyrus | L | V1 | -3 | -82 | -10 | 3.35 |  |
| Occipitotemporal cortex | R | PIT | 39 | -91 | -8 | 4.64 | 1585 |
| - Lateral occipitotemporal cortex | R | V4t | 49 | -72 | 1 | 4.26 |  |
| - Occipital pole | R | V3 | 36 | -93 | 0 | 4.12 |  |
| - Lateral occipital cortex | R | LO2 | 48 | -82 | -2 | 4.01 |  |
| - Posterior inferior temporal cortex | R | PIT | 45 | -83 | -9 | 3.86 |  |
| Visual cortex | R | V1* | 18 | -97 | 4 | 4.89 | 926 |
| - Occipital pole | R | V2 | 23 | -101 | 10 | 3.97 |  |
| **Subcortical** |  |  |  |  |  |  |  |
| Cerebellum Crus I | R |  | 46 | -63 | -44 | 5.96 | 2753 |
| - Cerebellum Crus II | R |  | 47 | -59 | -47 | 5.79 |  |
| Hipocampus cornu ammonis | L | Hipp | -27 | -22 | -21 | 5.73 | 2199 |

^*Juelich Histological Atlas^

**Supplementary Table 4** Whole-brain *t*-test results for Mode (share > private) effects, shown separately for each group (*Z* ≥ 3.1; *P* = 0.05, cluster-corrected, grey matter masked)

| **Region** |  | **HCPex** | **x** | **y** | **z** | ***Z*** | **Cluster size** |
| --- | --- | --- | --- | --- | --- | --- | --- |
| *AWNS* |  |  |  |  |  |  |  |
| **Frontal** |  |  |  |  |  |  |  |
| Medial prefrontal cortex | L | 9m | -7 | 55 | 16 | 5.21 | 13285 |
| - Ventral frontal pole | L | 10v | -3 | 58 | -15 | 5.04 |  |
| Superior frontal language area | R | SFL | 10 | 18 | 65 | 4.34 | 2482 |
| - Supplementary and cingulate eye field | R | SCEF | 4 | -2 | 68 | 4.32 |  |
| - Medial premotor cortex | R | 6ma | 12 | 14 | 64 | 4.04 |  |
| Medial prefrontal cortex | L | 8BM | -6 | 38 | 47 | 4.31 | 567 |
| - Medial prefrontal cortex | L | 9m | -4 | 45 | 44 | 3.66 |  |
| **Parietal & Occipital** |  |  |  |  |  |  |  |
| Dorsal posterior cingulate cortex | L | d23ab | -1 | -51 | 20 | 4.03 | 1109 |
| - Ventral posterior cingulate cortex | L | v23ab | -5 | -54 | 18 | 3.94 |  |
| - Posterior cingulate–precuneus region | L | 31pd | -3 | -58 | 31 | 3.70 |  |
| - Parieto-occipital sulcus | L | POS1 | -7 | -60 | 15 | 3.16 |  |
| **Subcortical** |  |  |  |  |  |  |  |
| Caudate nucleus | L | Caud | -9 | 10 | 3 | 5.55 | 2012 |
| Ventral anterior thalamic nucleus | L | VA | -11 | -2 | 10 | 4.61 |  |
| Caudate Nucleus | R | Caud | 8 | 11 | -4 | 4.72 | 1777 |
| Mediodorsal thalamus | R | Mdm | 3 | -8 | 5 | 3.9 |  |
| Cerebellum, Crus I | R |  | 33 | -89 | -33 | 4.21 | 934 |
| - Cerebellum, Crus II | R |  | 26 | -89 | -36 | 3.48 |  |
| Cerebellum, Crus II | L |  | -32 | -84 | -38 | 4.31 | 687 |
| - Cerebellum, Crus I | L |  | -32 | -86 | -35 | 4.23 |  |
| *AWS* |  |  |  |  |  |  |  |
| **Frontal** |  |  |  |  |  |  |  |
| Medial prefrontal cortex | L | 9m | -5 | 54 | 23 | 6.11 | 20345 |
| - Medial superior frontal cortex | L | 8BM | -3 | 19 | 54 | 5.32 |  |
| - Anterior dorsolateral prefrontal cortex | L | 9a | -11 | 56 | 31 | 4.70 |  |
| Inferior frontal gyrus, pars orbitalis | L | 47l | -43 | 28 | -6 | 4.93 | 6245 |
| - Inferior frontal gyrus, pars orbitalis, sulcal portion | L | 47s | -44 | 22 | -19 | 4.83 |  |
| Anterior cingulate cortex | L |  | -2 | 56 | -14 | 5.42 | 4200 |
| - Orbitofrontal cortex | L | OFC | -7 | 42 | -22 | 4.21 |  |
| - Ventral frontal pole | L | 10v | -3 | 41 | -22 | 4.15 |  |
| - Orbitofrontal cortex | R | OFC | 1 | 44 | -22 | 4.13 |  |
| - Rostral frontal pole | L | 10r | -10 | 49 | -15 | 3.94 |  |
| Anterior cingulate cortex | R | a24pr | 4 | 30 | 27 | 4.21 | 811 |
| - Pregenual anterior cingulate cortex | R | 33pr | 7 | 25 | 20 | 3.72 |  |
| **Parietal & Occipital** |  |  |  |  |  |  |  |
| Dorsal posterior cingulate cortex | L | d23ab | -1 | -53 | 26 | 5.88 | 5071 |
| - Medial superior parietal cortex | L | 7m | -5 | -64 | 26 | 4.44 |  |
| - Ventral posterior cingulate cortex | L | v23ab | -4 | -52 | 15 | 4.15 |  |
| - Ventral posterior medial parietal cortex | L | 31pv | -11 | -51 | 30 | 4.13 |  |
| - Posterior cingulate–precuneus region | L | 31pd | -10 | -51 | 32 | 4.01 |  |
| **Temporal** |  |  |  |  |  |  |  |
| Superior temporal auditory cortex | L | TE1a | -66 | -12 | -18 | 5.03 | 4361 |
| - Posterior superior temporal auditory cortex | L | TE1p | -63 | -35 | -1 | 4.85 |  |
| - Ventral posterior superior temporal sulcus | L | STSvp | -61 | -38 | 1 | 4.58 |  |
| **Subcortical** |  |  |  |  |  |  |  |
| Caudate nucleus | L | Caud | -7 | 9 | -4 | 5.30 | 3503 |
| - Nucleus accumbens | L | NAc | -8 | 7 | -10 | 4.96 |  |
| Caudate nucleus | R | Caud | 8 | 11 | -3 | 5.25 | 3035 |
| - Subgenual anterior cingulate cortex | R | 25 | 7 | 4 | -10 | 4.63 |  |
| - Ventral anterior thalamic nucleus | R | VA | 12 | -1 | 12 | 4.33 |  |
| *AWNS > AWS* none | | | | | | | |
| *AWS > AWNS* none | | | | | | | |

**Supplementary Table 5** Whole-brain *t*-test results for Person (self > other) effects, shown separately for each group (*Z* ≥ 3.1; *P* = 0.05, cluster-corrected, grey matter masked)

| **Region** |  | **HCPex** | **x** | **y** | **z** | ***Z*** | **Cluster size** |
| --- | --- | --- | --- | --- | --- | --- | --- |
| *AWNS* |  |  |  |  |  |  |  |
| **Frontal** |  |  |  |  |  |  |  |
| Anterior cingulate cortex | L | a24 | -6 | 37 | -6 | 7.87 | 28213 |
| - Anterior cingulate cortex | R |  | 6 | 39 | -9 | 7.35 |  |
| - Anterior cingulate cortex | L | p24 | -3 | 37 | 3 | 6.76 |  |
| - Anterior cingulate cortex | L |  | -2 | 33 | 3 | 6.61 |  |
| - Nucleus accumbens | L | NAc | -7 | 6 | -9 | 6.18 |  |
| - Anterior cingulate cortex | L | p24a | -3 | 40 | 9 | 5.98 |  |
| Frontal orbital cortex | L | 13l | -23 | 32 | -20 | 5.50 | 2910 |
| Inferior frontal gyrus, pars orbitalis | L | p47r | -44 | 39 | 5 | 4.80 | 1049 |
| Inferior frontal junction | L | IFja | -49 | 10 | 20 | 4.33 | 815 |
| Orbitofrontal cortex | R | OFC | 14 | 40 | -25 | 3.86 | 143 |
| **Temporal** |  |  |  |  |  |  |  |
| Posterior inferior temporal gyrus | L | TE2p | -49 | -45 | -16 | 5.21 | 3798 |
| Hippocampus | L | Hipp | -23 | -25 | -15 | 4.47 | 1298 |
| **Parietal & Occipital** |  |  |  |  |  |  |  |
| Parieto-occipital sulcus | L | POS1 | -16 | -56 | 9 | 4.26 | 753 |
| Occipital pole | R | V2 | 24 | -100 | 8 | 3.97 | 643 |
| **Subcortical** |  |  |  |  |  |  |  |
| Brainstem extending into the ventral tegmental area and dorsal raphe nucleus | L |  | -2 | -19 | -18 | 4.37 | 1017 |
| *AWS* |  |  |  |  |  |  |  |
| **Frontal** |  |  |  |  |  |  |  |
| Anterior cingulate cortex | L | p24 | -4 | 44 | 4 | 6.18 | 12749 |
| - Anterior cingulate cortex | L | a24 | -1 | 41 | 10 | 5.79 |  |
| Frontal pole, posterior portion | L | p10p | -23 | 68 | 5 | 4.34 | 882 |
| - Frontal pole, dorsal portion | L | 10d | -9 | 70 | 10 | 3.58 |  |
| **Parietal & Occipital** |  |  |  |  |  |  |  |
| Parieto-occipital sulcus | L | POS1 | -13 | -62 | 8 | 4.16 | 668 |
| **Subcortical** |  |  |  |  |  |  |  |
| Cerebellum, Crus II | R |  | 45 | -64 | -45 | 5.21 | 2485 |
| - Cerebellum, Crus I | R |  | 48 | -62 | -42 | 5.13 |  |
| *AWNS > AWS* |  |  |  |  |  |  |  |
| **Frontal** |  |  |  |  |  |  |  |
| Anterior cingulate cortex | L | a24 | -6 | 33 | -4 | 4.03 | 408 |
| *AWS > AWNS* |  |  |  |  |  |  |  |
| **Parietal** |  |  |  |  |  |  |  |
| Medial inferior parietal cortex | L | PFm | -57 | -61 | 33 | 3.95 | 1001 |

**Supplementary Table 6** Cluster-level correlation with Premonitory Awareness in Stuttering (PAiS, *Z* < 3.1, *P* < 0.05)

| **Region** |  | **HCPex** | **x** | **y** | **z** | ***Z*** | **Cluster size** |
| --- | --- | --- | --- | --- | --- | --- | --- |
| *Positive correlation: share > private* |  |  |  |  |  |  |  |
| **Subcortical** |  |  |  |  |  |  |  |
| Nucleus accumbens | R | NAc | 10 | 10 | -11 | 4.80 | 421 |
| *Negative correlation: share > private* | |  |  |  |  |  |  |
| **Frontal** |  |  |  |  |  |  |  |
| Inferior frontal cortex, pars opercularis | L | 44 | -50 | 18 | 5 | 5.3 | 668 |
| - Inferior frontal gyrus, pars orbitalis | L | 47l | -46 | 23 | -6 | 3.84 |  |
| - Inferior frontal gyrus, pars triangularis | L | 45 | -48 | 21 | -3 | 3.60 |  |
| **Temporal** |  |  |  |  |  |  |  |
| Ventral posterior superior temporal sulcus | L | STSvp | -60 | -20 | -11 | 4.10 | 327 |
| - Posterior superior temporal auditory cortex | L | TE1p | -66 | -32 | -4 | 4.01 |  |
| - Middle temporal gyrus | L | TE1m | -64 | -30 | -9 | 3.94 |  |
| *Positive correlation: self > other* none | | | | | | | |
| *Negative correlation: self > other* none | | | | | | | |

**Supplementary Table 7** Partial correlations between PAiS scores and share > private brain activity, controlling for trait anxiety and professional education (Bonferroni alpha = 0.05/3 ≈ 0.017). PAiS showed no statistically significant correlation with trait anxiety (*r* = -0.03, *p* = 0.856) or professional education (*r* = 0.16, *p* = 0.355).

|  | Right NAc | Left STS | Left IFG (BA 44,45,47) |
| --- | --- | --- | --- |
| PAiS | **0.55 (<.001)** | **-.52 (.002)** | -.36 (.041) |
| STADI trait anxiety* | .03 (.887) | .001 (.994) | .16 (.387) |
| Prof. education | -.13 (.464) | -.06 (.729) | .04 (.808) |

*Percentile rank

**Supplementary Table 8** Cluster-level correlation with Overall Assessment of the Speaker’s Experience of Stuttering (OASES, *Z* < 3.1, *P* < 0.05)

| **Region** |  | **HCPex** | **x** | **y** | **z** | ***Z*** | **Cluster size** |
| --- | --- | --- | --- | --- | --- | --- | --- |
| *Positive correlation: share > private* |  |  |  |  |  |  |  |
| **Frontal** |  |  |  |  |  |  |  |
| Supplementary and cingulate eye field |  | SCEF | 0 | 9 | 56 | 5.35 | 2154 |
| - Medial superior frontal cortex | L | 8BM | -4 | 26 | 50 | 5.07 |  |
| - Medial superior frontal cortex | R | 8BM | 6 | 20 | 56 | 5.01 |  |
| Medial prefrontal cortex |  | 9m | -13 | 56 | 21 | 5.14 | 1426 |
| - Posterior dorsolateral prefrontal cortex | L | 9p | -16 | 45 | 35 | 4.46 |  |
| - Anterior cingulate cortex | L | p24 | -11 | 41 | 7 | 4.41 |  |
| Inferior frontal gyrus, pars orbitalis, sulcal portion | L | 47s | -32 | 18 | -19 | 5.14 | 666 |
| Inferior frontal gyrus, pars triangularis | L | 45 | -46 | 21 | -2 | 5.43 | 653 |
| - Inferior frontal gyrus, pars orbitalis | L | 47l | -43 | 23 | -9 | 4.40 |  |
| - Frontal operculum | L | FOP4 | -53 | 16 | 2 | 5.94 |  |
| Anterior cingulate cortex | L | d32 | -5 | 42 | 24 | 4.30 | 378 |
| - Anterior cingulate cortex | L | p24 | -6 | 37 | 17 | 3.78 |  |
| Medial prefrontal cortex | R | 9m | 11 | 52 | 27 | 4.61 | 288 |
| - Medial prefrontal cortex | R | 9m | 6 | 56 | 23 | 3.91 |  |
| *Negative correlation: share > private* none | | | | | | | |
| *Positive correlation: self > other* none | | | | | | | |
| *Negative correlation: self > other* none | | | | | | | |

**Supplementary Table 9** Partial correlations between the Overall Assessment of the Speaker’s Experience of Stuttering (OASES) scores and share > private brain activity, controlling for trait anxiety and professional education (Bonferroni alpha = 0.05/6 ≈ 0.008). OASES total score and trait anxiety are highly positively correlated with *r* = 0.72, *p* < 0.001, while professional correlation showed a trending correlation with OASES with *r* = -0.32, *p* = 0.069.

|  | 8 BM SCEF | Left 9m/p24 | Left BA 47 | Left BA 45 | Right 9m | Left d32 |
| --- | --- | --- | --- | --- | --- | --- |
| OASES | **.56 (<.001)** | **.54 (.002)** | **.47 (.007)** | .45 (.010) | **.52 (.002)** | .41 (.019) |
| STADI trait anxiety* | -.21 (.257) | -.23 (.21) | -.13 (.473) | -.18 (.336) | -.24 (.179) | -.03 (.880) |
| Prof. education | -.05 (.784) | .02 (.909) | -.23 (.198) | -.04 (.811) | -.24 (.195) | .06 (.762) |

*Percentile rank

**Stimulus material**

Stimulus list for the behavioral experiment. Depending on the condition the question read for example “*Mag ich Abenteuer?”* (English: *Do I like adventures?*) or *“Mag Merkel Abenteuer?”* (English: *Does Merkel like adventures?*).

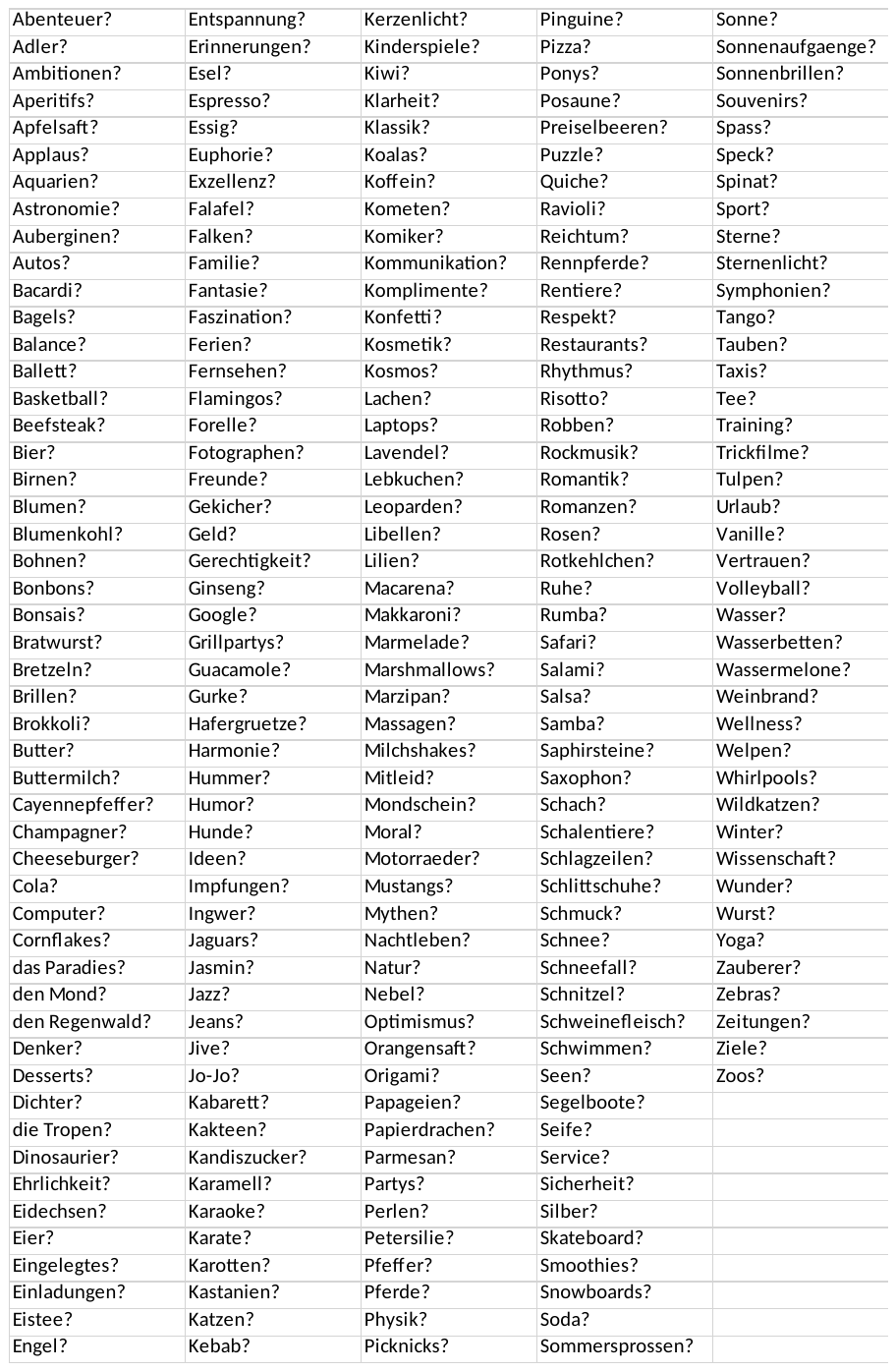

Stimulus list for the fMRI experiment. Depending on the condition the sentence read for example *“Ich mag Affen?”* (English: *I like monkeys?*) or *“Merkel* mag Amaretto*?”* (English: *Merkel likes amaretto?*).

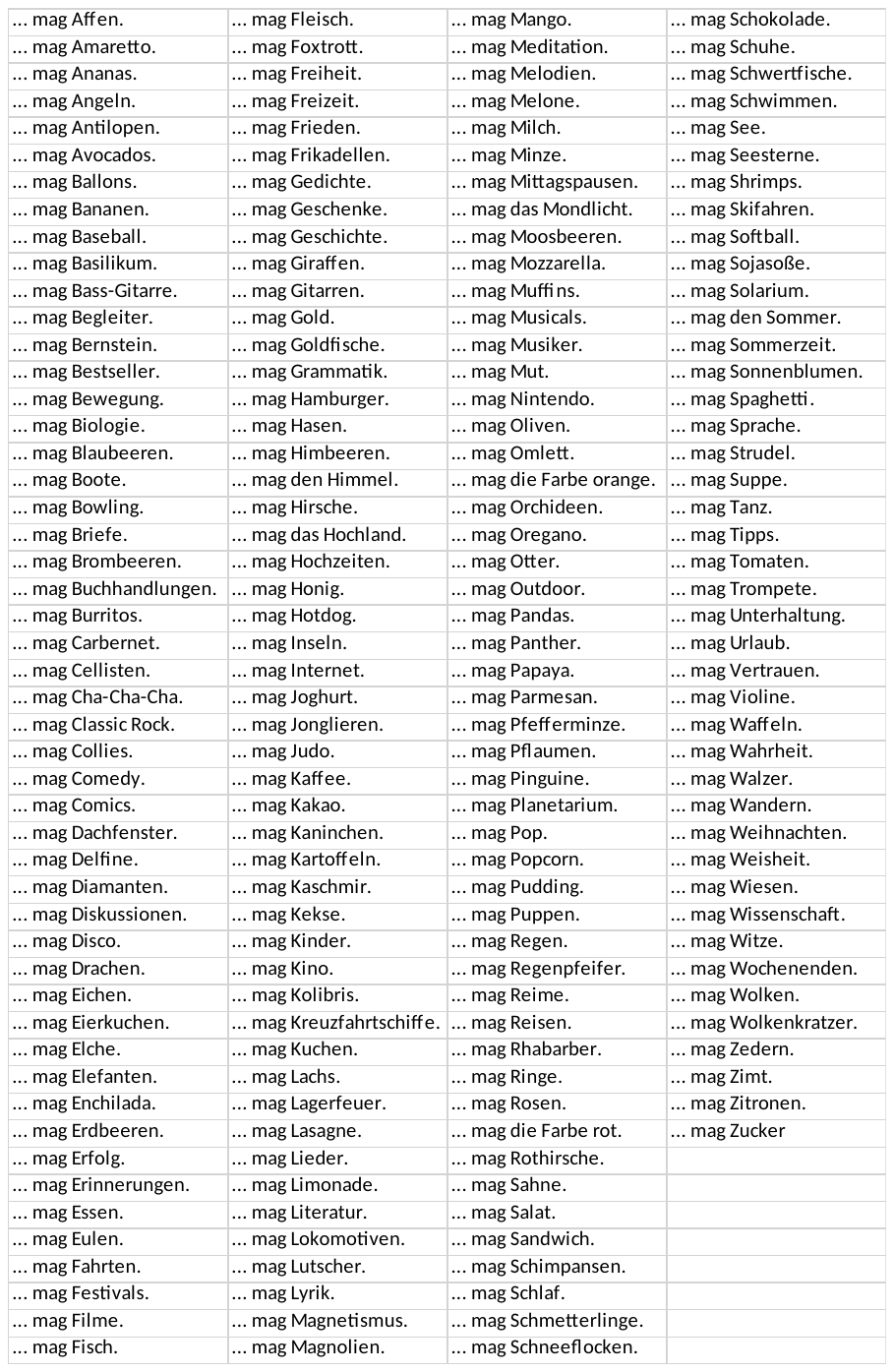
